# Genetic basis of aposematic coloration in a mimetic radiation of poison frogs

**DOI:** 10.1101/2023.04.20.537757

**Authors:** Tyler Linderoth, Diana Aguilar-Gómez, Emily White, Evan Twomey, Adam Stuckert, Ke Bi, Amy Ko, Natalie Graham, Joana L. Rocha, Jason Chang, Matthew D. MacManes, Kyle Summers, Rasmus Nielsen

## Abstract

The evolution of mimicry in a single species or population has rippling inter and intraspecific effects across ecological communities, providing a fascinating mechanism of phenotypic diversification. In this study we present the first identification of genes underlying Müllerian mimicry in a vertebrate, the Peruvian mimic poison frog, *Ranitomeya imitator*. We sequenced 124 *R. imitator* exomes and discovered loci with both strong divergence between different mimetic morphs and phenotypic associations within an intraspecific admixture zone, implicating *mc1r*, *asip*, *bsn*, *retsat*, and *krt8.2* in the evolution of mimetic color phenotypes. We confirmed these associations for most candidate genes through linkage mapping in a lab-reared pedigree. We also sequenced transcriptomes from the model species, allowing tests for introgression and revealing that the mimetic resemblance between *R. imitator* and the models evolved independently. Selection analyses of the candidate genes show that the mimicry phenotypes likely have evolved through selective sweeps acting on polygenic variation. Our results suggest that the evolutionary origins and molecular mechanisms underlying mimicry phenotypes in vertebrates may be radically different from those previously documented in invertebrates such as the iconic *Heliconius* butterfly mimicry complex.

**One Sentence Summary:** Müllerian mimicry evolved through independent selective sweeps on color and pattern loci in the mimic poison frog.

## Main Text

Mimetic radiations, in which different populations of a species diverge to resemble multiple models or co-mimics, have provided compelling examples of adaptation since the work of Wallace and Müeller (*1*). Butterflies of the genus *Heliconius* have been the main focus of genomic analyses of mimetic radiations, illuminating the genes and gene pathways controlling the color and pattern variation involved in mimicry (*2–4*). With a few exceptions (e.g. (*5*)), much of the resemblance between mimics and co-mimics in these radiations appears to stem from the exchange of mimicry genes through introgression (*6–8*), but it is unclear if this conclusion extends to other species, in particular vertebrates. In fact, there is no vertebrate system for which the molecular mechanism of mimicry has been determined. However, there is a vertebrate that shows remarkable parallels to the *Heliconius* mimicry system: the Peruvian mimic poison frog (*Ranitomeya imitator*) has undergone a mimetic radiation to resemble several existing morphs of congeners (i.e., model species) across northern Peru. We focused on a banded morph that mimics *R. summersi* and a striped morph that mimics *R. variabilis* (Fig. 1) (*9, 10*).

**Fig. 1.**
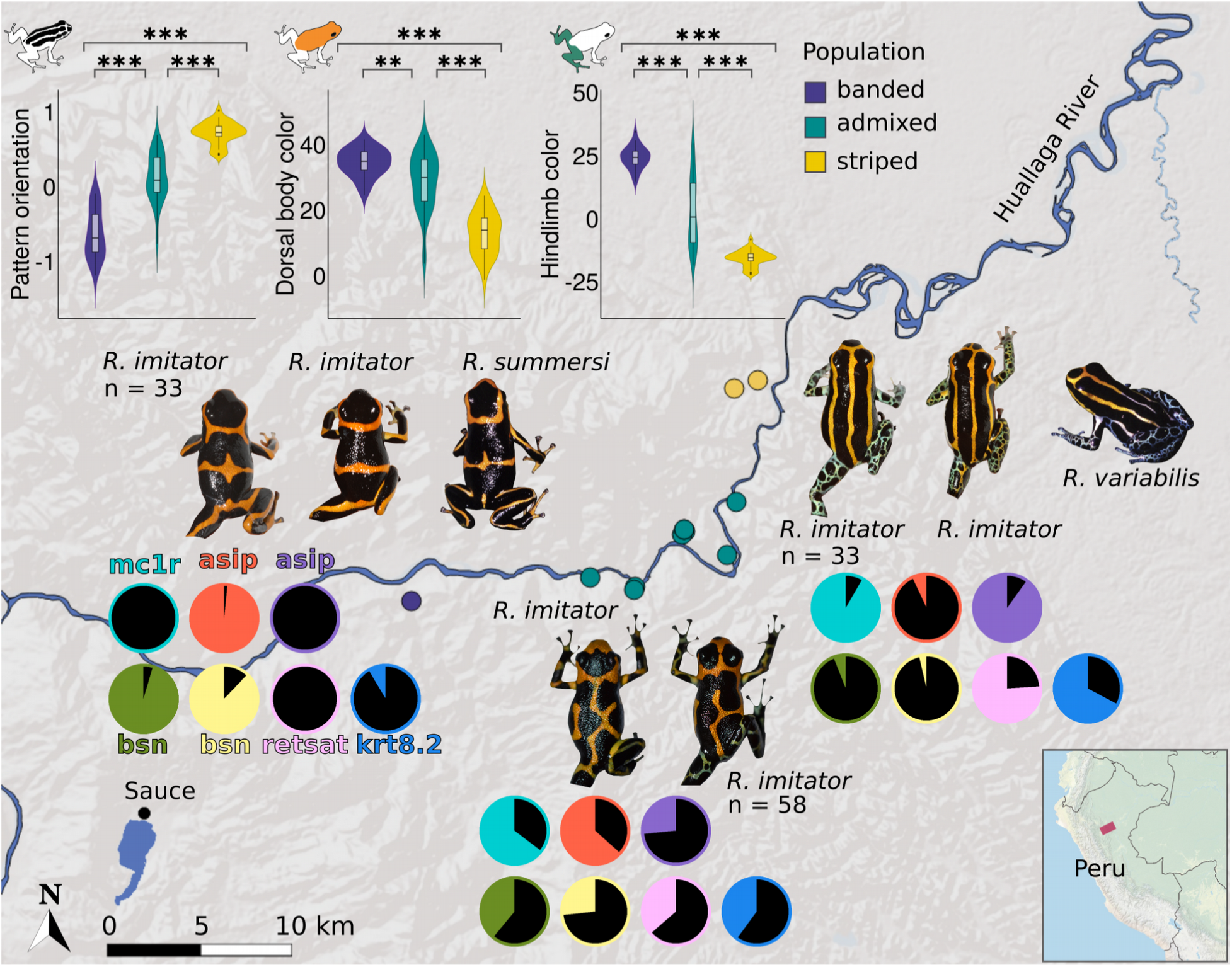
*Ranitomeya imitator* phenotypes and allele frequencies at candidate color genes. Map showing the sampling localities (small colored dots, red rectangle in the inset map) and sizes of the three main morphs of *R. imitator* in San Martin province, Peru. Traversing northeast along the Huallaga river region, *R. imitator* switches from a banded morph resembling sympatric *R. summersi* to a striped morph that mimics sympatric *R. variabilis* after passing through a 7 km-wide transition zone where *R. imitator* exhibits a spectrum of intermediate phenotypes. Differences in coloration across this mimicry gradient are significant as shown by the violin plots (two-tailed p-value for Mann-Whitney U: ** p < 0.01, *** p < 0.001). Pie charts depict the frequencies of reference (black) and alternate (non-black) alleles of different candidate gene SNPs that have the strongest association with at least one of the coloration phenotypes.

Amphibians have notoriously large, complex genomes, and poison frogs are no exception (*11*), making genome sequencing and assembly extremely challenging. To overcome this, we used a combination of exome capture, divergence mapping, association testing within admixed populations, and pedigree linkage analyses to identify genes underlying the divergence of the mimetic morphs of *R. imitator*. We then used population genomic analyses of transcriptome sequence data to determine whether the resemblance between the mimetic populations of *R. imitator* and their model species is due to introgression or independent evolution. Finally, we identified patterns of selection at the molecular level to elucidate the mechanisms of molecular adaptation.

## Population genomic characterization of the *Ranitomeya* complex

To enable population-scale genomic analyses in *Ranitomeya imitator*, which has an estimated genome size of 6.8 Gb (*12*), we captured and sequenced exons from 13,084 genes in 124 individuals representing three distinct phenotypes (Fig. 1). We also sequenced transcriptomes from the models *R. summersi* and *R. variabilis*. We mapped reads to a *de novo* exome assembly and identified 928,822 high-quality genetic variants across the *Ranitomeya* complex. Most of this genetic variation is attributable to differences between *R. imitator* and the model species, *R. summersi* and *R. variabilis*, represented by PC1 in the principal component analysis (PCA) of exome-wide genetic variation (Fig. 2A, left panel). The tight cluster of *R. imitator* in the middle of PC2, which represents differences from the model species, hints that *R. imitator*’s phenotypic likeness to the models may not derive from introgression.

**Fig. 2.**
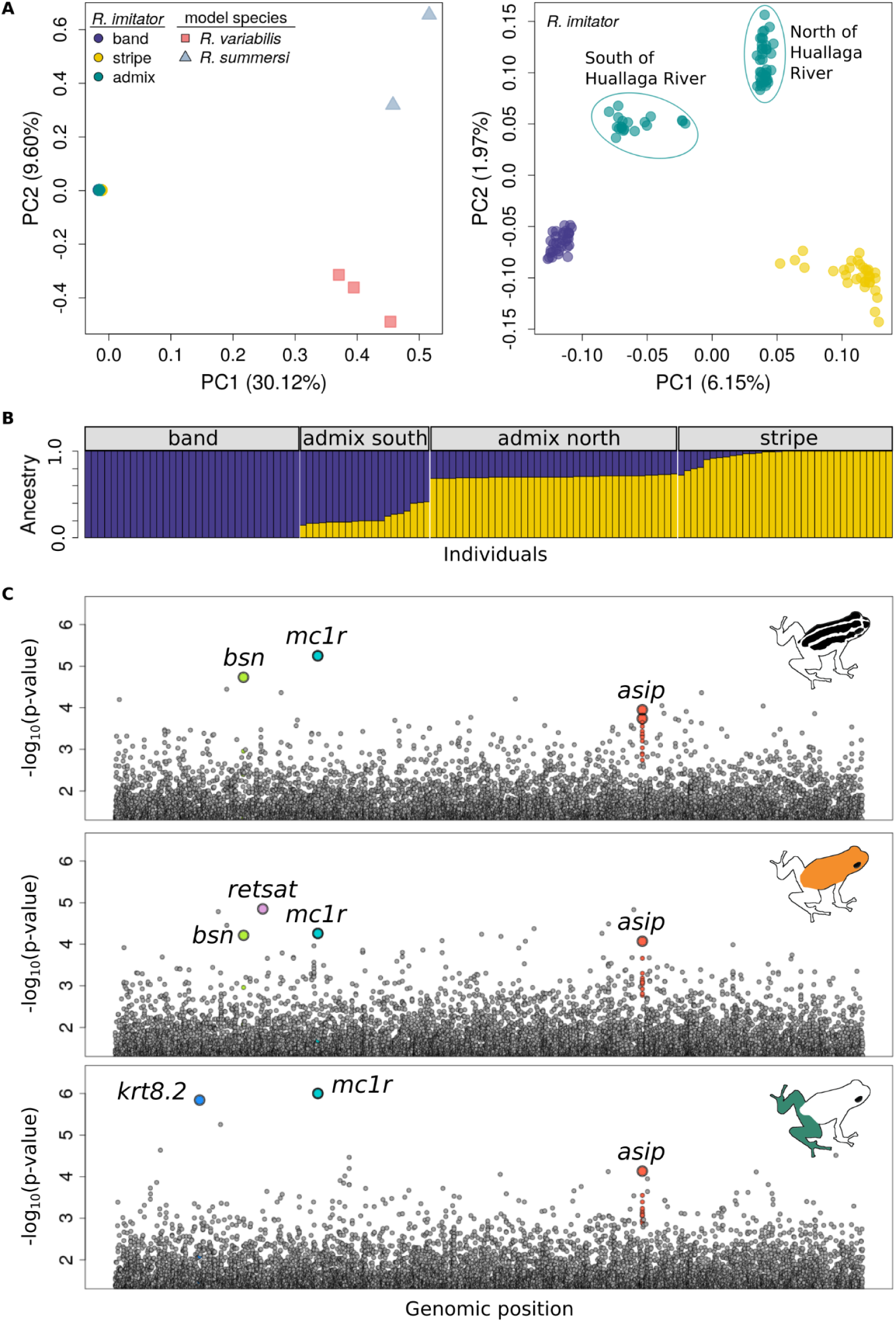
Genome-wide relationships among *Ranitomeya* and scans for genes underlying mimetic phenotypes. (**A**) Left panel: Principal component analysis (PCA) of exome-wide genetic variation among *R. imitator*, *R. summersi* (banded), and *R. variabilis* (striped) shows clear divergence between *R. imitator* and the models. Right panel: PCA of the genetic variation within *R. imitator* places individuals from the introgression zone between the striped and banded populations along PC1. The second axis of genetic variation, PC2, distinguishes the admixed frogs from the striped and banded populations. (**B**) NGSadmix (*57*) analysis showing different intermediate proportions of striped versus banded ancestry (colored bars) among admixed frogs sampled north and south of the Huallaga River. (**C**) Manhattan plots of -log_10_-scaled Fisher’s combined p-values from the test for excessive allele frequency differences between banded versus striped morph populations and tests of association between genotypes of admixed individuals and different aspects of coloration. Scans for SNPs associated with pattern orientation, dorsal body color, and hindlimb color are shown in the top, middle, and bottom panels, respectively. Exons are arbitrarily ordered along the x-axes. SNPs (points) in candidate coloration genes are displayed in color.

Among 295,279 SNPs that segregate within *R. imitator,* most genetic differences are between the banded and striped morph populations, defining PC1 of *R. imitator* variation (Fig. 2A, right panel). The exome-wide F_ST_ between the striped and banded morphs is 0.17, which is appropriate for identification of candidate regions underlying color differences through scans of divergence (*13*). The admixed individuals are positioned between the pure morphs on PC1, confirming that this population is genetically intermediate (*10*), and are further separated into two clusters coinciding with subpopulations north and south of the Huallaga River (Fig. 1). There is greater genetic similarity among admixed individuals to the pure morph population from the same side of the river, suggesting that the river is a barrier to gene flow that favored higher migration from the morph residing on the like side. This is consistent with relative amounts of striped/banded ancestry inferred in admixture analyses (Fig. 2B). Importantly for the purposes of trait mapping, the levels of ancestry are fairly uniform across individuals within the north-of-river and south-of-river admixed subpopulations (Fig. 2B), suggesting that the admixture zone is sufficiently old (*14*) for recombination to break up genetic linkage. This is also consistent with the observation that admixed individuals are separated from the pure morph populations along PC2 of the PCA (Fig. 2A, right panel).

## Genetic basis of mimetic phenotypes

### Divergence mapping

We performed a scan across 266,944 SNPs distributed in 12,972 genes in 33 pure striped and 33 banded *R. imitator* to identify loci with higher allele frequency differences (*δ*) than expected given the background divergence between these two morphs. We assigned approximate p-values to outliers using simulations based on the Balding-Nichols model (*15, 16*) for the joint distribution of allele frequencies between the two morphs. Even very divergent SNPs (several with *δ* > 90%) were not significant after Bonferroni corrections to account for the number of tested SNPs. However, these corrections are very conservative and among the 20 most divergent SNPs (table S1) across the exome (> 99 *δ* percentile) we identified 11 SNPs in three genes (*mc1r*, *asip*, *bsn*) with relevant functions for influencing color phenotypes.

The most diverged SNP in the *R. imitator* exome (*δ* = 92%) was located in the melanocortin 1 receptor gene, *mc1r*, which encodes a protein that operates in the synthesis of melanin pigments and melanosome aggregation. This gene has been extensively documented as affecting pigmentation in various taxa (*17–20*). The next most diverged SNP (*δ* = 91%) is located in the agouti-signaling protein gene, *asip*, which produces a protein that competes with melanocortins to bind Mc1r, thereby inhibiting melanin dispersion in melanocytes. Increased *asip* expression has been linked to lighter coat colors in mammals (*21, 22*) and the countershading patterning of fish (*23*). Among the 20 most divergent SNPs across the exome, nearly half (eight) were located in *asip*. A third candidate gene, *bsn*, harbors the seventh most diverged SNP (*δ* = 89%). This gene has not previously been implicated in color phenotypes, but its product, Bassoon, is a scaffolding protein which could play a role in spatially organizing organelles involved in pigmentation such as melanosomes within chromatophores.

### Dissecting the genetic basis for different features of mimetic phenotypes

Next, we tested for associations between 160,285 SNPs from 12,739 genes and quantitative measures of pattern orientation, dorsal body color, and hindlimb color in 58 admixed individuals from an introgression zone between the striped and banded morphs of *R. imitator* (Fig. 1). We combined the p-values from this association test with those of the divergence test using Fisher’s Method (*24*) since SNPs with the largest combined test statistic, *X*^2^, are both highly divergent between the striped and banded morphs and highly correlated with a continuous color feature in admixed individuals, consistent with a causal color locus. Though no SNPs remained significant after a Bonferonni correction, all three candidate loci from the divergence mapping (*mc1r*, *asip*, *bsn*) were among the top 20 (> 99.9 percentile) highest *X*^2^ values for either pattern orientation, dorsal body color, and/or hindlimb color (Fig. 2C, tables S5-7).

Combining evidence across mapping experiments also revealed two additional strong candidate color genes, *retsat* and *krt8.2*. *Retsat,* the highest-ranked *X*^2^ SNP for dorsal color (table S6), is involved in retinol metabolism which could influence the orange hue of the dorsum through regulation of carotenoid levels. *Krt8.2*, the third-ranking *X*^2^ SNP for hindlimb color, which encodes a type II basic intermediate filament protein that polymerizes with other keratins to form a cytoskeletal component of epithelial cells, could play a role in generating the blue/green hues of the hindlimbs since keratins are involved in structurally generating these colors in birds (*25, 26*).

SNPs with the largest *X*^2^ values (strongest association) in candidate genes for each respective coloration phenotype on average explained 7%, 6%, and 9% of the pattern, dorsal color, and hindlimb color variation, respectively, in the admixed frogs (table S8, see “Phenotypic variance explained by candidate SNPs” of supplementary text), with the amount of phenotypic variance explained by any single candidate SNP ranging from 1.2-17.6%.

### Linkage analyses in pedigrees

To validate the candidate genes identified by divergence and association mapping, we performed linkage analyses in a captive pedigree of seven *R. imitator* families. Inter-morph crosses between three banded and three striped parents produced seven families with nine F2 individuals on average (1-15 F2s per family), totalling 14 F1 and 62 F2 individuals across the entire pedigree (Fig. 3A).

**Fig. 3.**
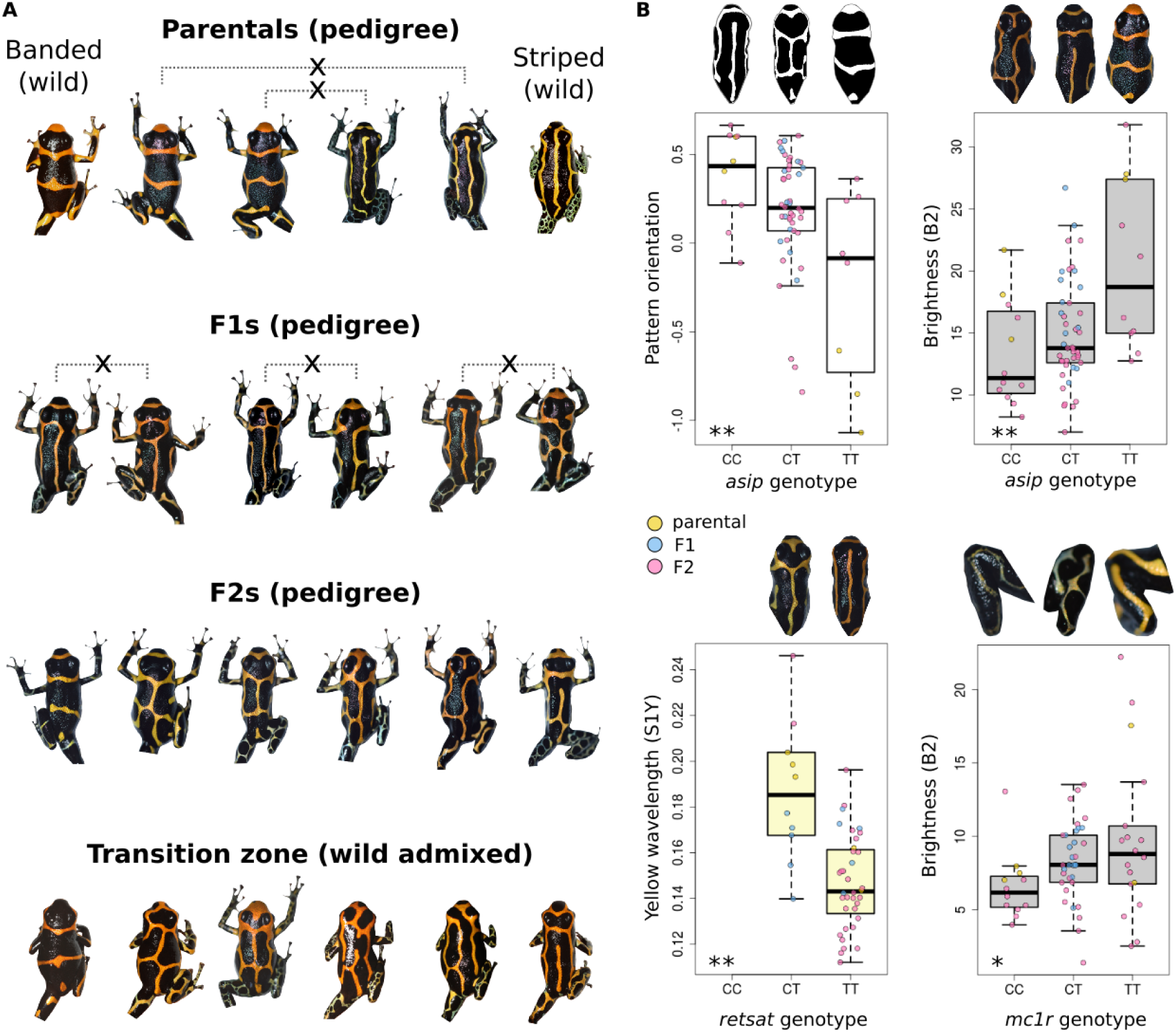
Relationship between candidate SNP genotypes and phenotypes in a captive pedigree of *R. imitator*. (**A**) Crosses initiated between captive striped and banded *R. imitator* produced F1 and subsequent F2 generation offspring exhibiting phenotypes that closely mirror those of wild frogs from the admixture zone. (**B**) Boxplots show significant (LOD score q-value < 0.1*, q-value < 0.01**) relationships between allelic dosage at candidate gene SNPs and different color or pattern phenotypes of *R. imitator* crossed in captivity for three generations. The FDR-corrected q-scores are calculated for a total of 14 tests involving one SNP from each of four candidate genes and five phenotypic features.

The spectrum of hues and patterns on the body and legs of the pedigree population closely resembled that of wild frogs (Fig. 3A). Each of the genotyped SNPs from the putative pattern candidate genes, *mc1r*, *asip*, and *bsn*, showed significant associations with pattern orientation among pedigree frogs (table S9, LOD range 0.93-2.54, q-value range 0.003 - 0.091). Of the candidate genes for dorsal body color, the genotyped *asip* SNP was significantly associated with body brightness (table S9, LOD = 2.47, q-value = 0.003) while *retsat* had a highly significant association with yellow intensity on the body (LOD = 3.49, q-value = 8.54×10^-4^). Neither *mc1r* or *bsn* were significantly associated with body brightness or yellow intensity in the pedigree (LOD scores ranging 0.09-0.48, q-values = 0.238-0.571). For the hindlimbs, we tested SNPs in the hindlimb color candidate genes *mc1r* and *asip* for associations with brightness and green wavelengths in the pedigree (table S9). *Mc1r* was significantly associated with hindlimb brightness (LOD = 1.13, q-value = 0.064), while *asip* was marginally so (LOD = 0.62, q-value = 0.182). See “Linkage analysis in a captive *R. imitator* pedigree” section of the supplement for further details.

## Mimetic phenotypes were not facilitated by introgression

In butterfly mimicry systems some species closely match model phenotypes due to alleles acquired through introgression from the models (*8, 27*). To determine whether this is the case for *R. imitator,* we tested for patterns of excess genome-wide allele sharing between *R. imitator* and their respective model species using ABBA-BABA tests (Fig. 4A, table S10), which provided no evidence for admixture between *R. imitator* and any of the models (D-statistic = −0.002, p-value = 0.6). Similarly, neighbor-joining trees for the specific genes associated with the mimetic phenotypes show clear separation by species (Fig. 4B), corroborating that there are no shared haplotypes between *R. imitator* and the model species.

**Fig. 4.**
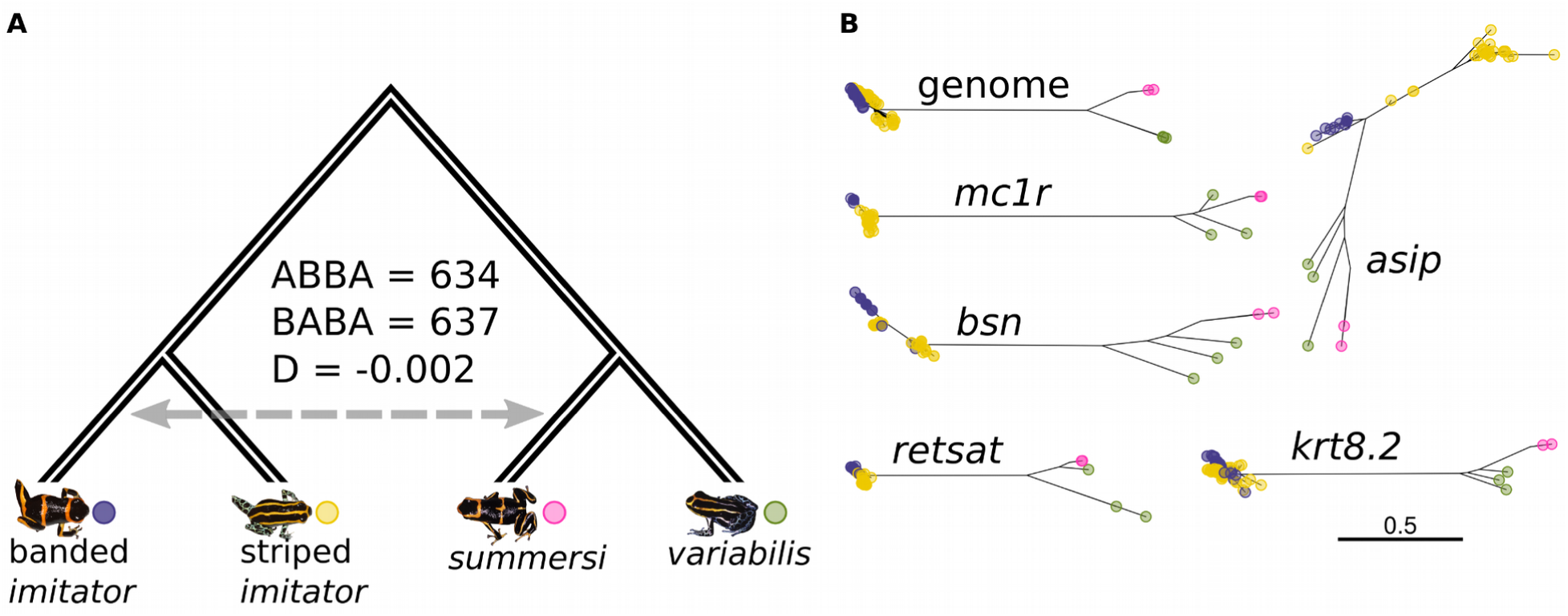
Genetic comparisons between different *R. imitator* morphs and the species that they mimic. **(a)** Genome-wide patterns of ABBA-BABA allele topologies among the banded and striped *R. imitator* populations and their respective model species, *R. summersi* (banded) and *R. variabilis* (striped), shows no enrichment of BABA patterns (D = −0.002, p-value = 0.6), suggesting no introgression between *R. imitator* and the models. **(b)** Unrooted neighbor-joining trees show that all banded (purple dots) and striped (yellow dots) *R. imitator* individuals are clearly more phylogenetically distant to *R. summersi* (pink dots) and *R. variabilis* (green dots) individuals than they are to each other, both across the entire genome and at each of the candidate coloration genes.

## Evolution of mimicry in *R. imitator*

We used Hudson–Kreitman–Aguadé (HKA) tests (*28*) to determine whether levels of polymorphism within *R. imitator* relative to divergence from the model species was consistent with neutral evolution at the candidate gene exons that contained the strongest phenotype-associated SNPs (table S11). Based on *X*^2^_1_ approximations with genomic control for the goodness-of-fit between observations and neutral expectations, these tests rejected neutrality at the candidate exons of *mc1r*, *bsn*, and *retsat* (*X*^2^_imitator_ = 5.82 - 18.35, q = 1.101e-4 - 0.032). Furthermore, there was significant evidence (*X*^2^= 4.46 - 22.44, q = 1.304e-5 - 0.046) of non-neutral evolution at candidate exons of *mc1r*, *asip*, *bsn*, and *retsat* within at least one of the pure *R. imitator* morphs (table S11). In all cases where exons significantly deviated from neutral expectations, we observed decreased polymorphism within species or populations compatible with selective sweeps (table S11). An alternative approach for obtaining p-values using coalescent simulations, instead of genomic control, also supported non-neutral evolution at the candidate genes (see “Tests of neutral evolution at candidate color genes” of the supplementary text).

## Discussion

Coloration is one of the most eco-evolutionarily important traits of dendrobatid frogs, yet little is known regarding its genetic basis (*29*), except for a few transcriptomic studies that generated extensive lists of candidate genes (*12, 30*). We used divergence and association mapping to identify candidate genes, which we subsequently validated using linkage analyses in pedigrees. While some of these candidate genes, particularly *mc1r* and *asip*, have been identified to influence color in many previous studies, we also discovered new genes such as *bsn* and *krt8.2*.

*Mc1r* and its competitive antagonist, *asip*, are directly involved in melanogenesis and have been linked to color phenotypes across taxa (*31–36*), including the harlequin poison frog, *Oophaga histrionica*, where *mc1r-*mediated melanosome density influences dorsal background color (*37*). In *R. imitator*, both *mc1r* and *asip* appear to be involved in pattern formation, dorsal body color, and hindlimb color. Another gene that appears to be involved in dorsal pattern and color phenotypes is *bsn*, which encodes a large scaffolding protein (Bassoon) involved in the structural and functional organization of the cytomatrix of the active zone in synapses (*38*). The iridophore platelets of dendrobatids are arranged in vertical stacks which require a mechanism to govern spacing and thickness, providing one possible role for such a scaffolding protein. Alternatively, it could be anchoring pigment-containing organelles (chromatophores) in specific fields of tissue to create bands, stripes, or other pattern elements. For example, in the retina Bassoon functions in anchoring the photoreceptor ribbon to the synapse (*39*). Additional studies are needed to understand the novel functional connection between *bsn* and color patterning.

The last two candidate genes, *retsat* and *krt8.2,* are exclusively associated with colors from different body parts. *Retsat* is associated with dorsal body color, a pattern found in previous studies on fish. *Retsatl* (a *retsat* homolog) is differentially expressed at higher levels in orange skin in clownfish (*40*) and yellow compared to white skin in an East African cichlid fish (*41*). Although the link between *retsat* and carotenoid coloration remains unclear, *retsat* is known to act on retinol, a carotenoid cleavage product (*42*), as well as galloxanthin, a colorless apocarotenoid (*43*). Given its link to yellow/orange coloration in other taxa and the range of reddish/orange to yellow coloration displayed on the dorsum of *R. imitator*, *retsat* is an intriguing color candidate gene whose function warrants further investigation. *Krt8.2*, which encodes a keratin protein found in epithelial cells, was associated with hindlimb color, which varies from orange to blue-green, the latter of which requires co-localization of melanosomes and overlying chromatophores (*25*). In another poison frog species, *Dendrobates auratus*, differential expression of this gene was associated with variation in skin color (*44*). So while keratin is not currently known to directly influence amphibian skin color, it is possible that keratinocytes may be playing a role in melanosome transport and organization (*45*) in poison frogs.

The identification of likely causal genes involved in *R. imitator* mimetic phenotypes provides the first opportunity for elucidating the evolution of mimicry in a vertebrate using molecular methods. Despite evidence that Müllerian mimicry operates in snakes (*46*), birds (*47*), catfish (*48*), bumblebees (*49*), ants (*50*), and beetles (*51*), virtually all of our empirical understanding of its genetic and evolutionary basis (*52*) stems from research on Neotropical butterflies (*53, 54*), (see (*50*) for an exception). This rich body of butterfly work has made it clear that species can evolve to mimic sympatric species through hybridization with the models, thereby obtaining mimetic variants through introgression. In contrast to these findings in butterflies, we find no evidence for introgression with the model species for any of the candidate genes in *R. imitator*. Instead, the large allele frequency differences between *R. imitator* morphs and reduced ratio of segregating to fixed differences with respect to the model species at the candidate genes, suggests rapid turnover of alleles at color and pattern loci driven by partial or full selective sweeps on standing or *de novo* variation. We might expect such a scenario in a dynamic ecological environment where the interaction of *R. imitator* populations with model species changes relatively rapidly on an evolutionary time scale.

The combined contribution of the candidate genes to the phenotypes is relatively low (sum of variance explained < 28%). This may partially be explained by measurement error and incomplete linkage disequilibrium between the most highly associated SNPs and causal variants, but also suggests that the mimicry phenotypes are somewhat polygenic. It should come as no surprise that such complex phenotypes have a complex genetic basis. However, in butterflies, complex mimicry color patterns have been found to be caused by a supergene controlling multiple phenotypes (*55*). Our observations of a polygenic basis, absence of introgression and evidence for partial local sweeps, suggests a quite different genetic and evolutionary model for the evolution of mimicry phenotypes in poison frogs and perhaps other vertebrates than that observed in butterflies.

In conclusion, this study provides, for the first time, a link between genetics and coloration mimicry in vertebrates, with candidate genes cross-validated through pedigree analyses. It appears that this mimicry evolved independently in the mimic poison frog (relative to the model species), likely through repeated partial or full sweeps, providing strong evidence for alternative routes to this type of diversification apart from introgression. This study also demonstrates a viable genomics-enabled path towards filling a stark gap in understanding the genetic basis of color phenotypes in amphibians (*56*) even in the absence of a reference genome.

## Acknowledgements

The authors wish to thank Pablo Venegas of CORBIDI for assistance with research, collection and export permits in Peru, and Manuel Panaifo of Chazuta, San Martin Province for assistance with fieldwork in Peru. We would also like to thank Lydia Smith at the Evolutionary Genomics Lab and Shana McDevitt at the QB3 Genomics core at UC Berkeley for their support in generating genomic data. Sarah Fitzpatrick’s lab and Chris Kozakiewicz provided helpful discussion in revising the manuscript. Tissue exports were authorized under Contrato de Acceso Marco a Recursos Geneticos 0009–2013-MINAGRI-DGFFS/DGEFFS, with CITES (Convention on International Trade in Endangered Species of Wild Fauna and Flora) permits number 003302 and 001718. Permission for the protocols used for research on vertebrate animals in this project was obtained from the ECU Institutional Animal Care and Use Committee (IACUC), under AUPs 281D and 376. Research Permits for collection and export of tissues were obtained from the Peruvian Servicio Forestal y de Fauna Silvestre (SERFOR): Authorizations 050– 2006-INRENA-IFFS-DCB, 067–2007-INRENA-IFFS-DCB, 005–2008-INRENA-IFFS-DCB, Resolución Directoral No. 137-2009-AG-DGFFS-DGEFFS, No. 0266-2011-AG-DGFFS-DGEFFS, No. 0402-2013-MINAGRI-DGFFS/DGEFFS and No. 232-2016-SERFOR/DGGSPFFS.

## Funding

National Science Foundation, in the form of a collaborative grant to K. Summers, R. Nielsen and M. MacManes from the Division of Environmental Biology, Evolutionary Processes: DEB1655336 (KS), DEB1655191 (RN), DEB1655585 (MM)

## Author Contributions

Conceptualization: TL, RN, KS

Methodology: TL, RN, KB

Formal Analysis: TL, DA, EW, AK, JLR, KB, MDM

Investigation: TL, EW, ET, AS, AK, NG, JC, MDM, KS

Resources: ET, AS, KS

Writing - original draft: TL, RN, KS, DA, EW

Writing - review and editing: TL, RN, KS, EW, ET, AS, AK, NG, JLR, JC, MDM

Visualization: TL

Supervision: RN, KS, TL

Project administration: RN, KS, TL

Funding acquisition: RN, KS, MDM

## Competing interests

Authors declare that they have no competing interests.

## Data and materials availability

All genetic sequencing and phenotypic data will be deposited to the Sequence Read Archive and Dryad. All code used in this study is available at https://github.com/tplinderoth/poison_frog_color/tree/main/R_imitator_color.

## Supplementary Materials

Materials and Methods

Supplementary Text

Figs. S1 to S10

Tables S1 to S12

References (58–99)

## Supplementary Materials for

### Materials and Methods

#### *Ranitomeya imitator* and model species samples

We sampled *R. imitator, R. variabilis and R. summersi* from sites throughout the San Martin province of Peru (Fig. 1) throughout the morning and early afternoon in 2010, 2012, 2013, and 2014. We hand-caught frogs as we encountered them out foraging during the day or resting in plant axils. Frogs were placed individually into film canisters or 15 mL Falcon tubes and transported back to a field-based laboratory where we collected phenotypic data and toe clips, which were stored in 96% ethanol. On the following day, frogs were released to the location from which they were caught. We collected samples from each location in one to three consecutive days. The original *R. imitator* parentals used to generate the multigenerational pedigree were obtained from a commercial breeder (Understory Enterprises LLC) that breeds different morphs of *R. imitator* from known localities in Peru.

#### Phenotyping

We quantified the color and pattern of *R. imitator* individuals from digital photographs taken with a Nikon D7000 camera using a Nikkor 85mm macro lens. Images were converted from RAW to the highest quality JPEG format using Nikon ViewNX2 software. Prior to phenotyping, we measured the anteroposterior length (running from the tip of the pelvis to the rostrum) of each frog as well as its midline orientation to the horizontal axis using imageJ v. 1.53c software (*58*). These measurements were used in R v. 3.6.3 (*59*) to standardize the size and orientation of the frog subjects across all of the photos. The standardized photos were imported into GIMP v. 2.8.22 (*60*), which we used to manually generate black masks for black portions of the frogs to reduce noise from glare and debris. We also used GIMP to isolate subjects from backgrounds and generate binary masks for restricting analyses to regions of interest (ROI). The preprocessed images were exported in PNG format for input into our phenotyping scripts. All scripts used for image processing and phenotyping are available at https://github.com/tplinderoth/image_phenotype.

We measured the proportion of the dorsal body surface (limbs excluded) that was black, denoted PB. This was calculated simply as the fraction of black pixels among all non-transparent pixels in the ROI. We also measured four aspects of color for the non-black areas in the CIELAB (L*a*b*) color space (*61*): dorsal body color, DC_a_, and hindlimb color, LC_a_, along the red-green color axis (a*), and dorsal body color, DC_L_, and hindlimb color, LC_L_, along the lightness axis (L*). To quantify color from PNG images we first determined the average red, green, and blue values in the RGB color space over all non-transparent pixels in the ROI. Specifically, for a raster image ROI with *d* non-transparent pixels having color values (*ψ_v,1_*,…,*ψ_v,d_*) for color channels *v∈* {red, green, blue }, the average for each RGB channel, 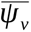, was calculated as

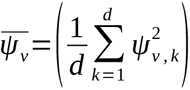

These average RGB values were then converted to L*a*b* color space using CIE Standard Illuminant D65 as the reference white for both color spaces.

Pattern was measured in terms of the degree to which black streaks run parallel to the midline axis along the body, denoted PT_rot_ and referred to as ‘pattern orientation’ throughout the text. To elaborate on how PT_rot_ is calculated from a raster image we impose a binary classification scheme for pixel color, which in our case is black and non-black. Let *γ_r_* and *γ_c_* denote the number of transitions between black and non-black pixels in a straight line along the entire horizontal axis and vertical axis of a raster image, respectively. Note that throughout we use the subscript *r* to refer to the horizontal axis of a raster image (the rows) and *c* to refer to the vertical axis (the columns). Pattern track length, *τ*, is defined as the number of directly adjacent black pixels along a straight line. Let *τ^max^_r_* denote the maximum *τ* in a set of tracks for a given raster image row, and similarly, *τ^max^_c_* denotes the maximum *τ* among the set of tracks for a given column. The effective segment length, *ϕ*, is the count of all pixels in a given row or column to be analyzed and so is bounded from above by the image’s width or height in pixels, respectively. For a raster image of width, *w*, and height, *h*

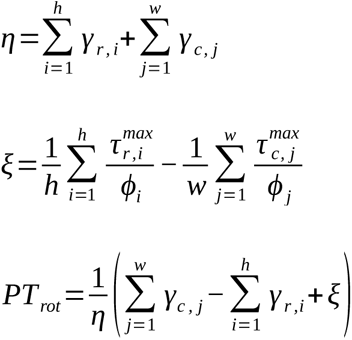

As PT_rot_ increases in the positive direction it corresponds to more stripedness, while increasing magnitude in the negative direction indicates more bandedness. A PT_rot_ value of 0 corresponds to a frog that is perfectly intermediate between striped and banded. We checked the effectivness of our methods for quantifying color and pattern by ensuring that image orderings using the rank values for each of the aformentioned measures were sensible (fig. S1-3).

We used an Ocean Optics USB spectrometer and the associated Ocean Optics software to collect spectral reflectance data for captive-bred frogs. We collected 10 data points (8 dorsal, 2 hindlimb, 3 ventral, see fig. S4) for each frog. We calibrated the spectrometer with a pure white standard between each frog to maintain accuracy of the measurements. We used the pavo package (*62*) in R to visualize, clean, and analyze the resulting spectral reflectance data. We first subdivided the data into groups with reference to body region, as shown in fig. S4. For each region (body and legs) on each frog, we aggregated the data points by taking the mean and set negative values to zero. Visualization showed noise at both ends of the spectral curve, which we controlled for by truncating the spectra to a wavelength range of 450 - 950 nm (previously 400 - 1041 nm). After processing the spectral reflectance data, we visualized them again, this time with the average and standard deviation plotted. We quantified mean brightness (B2), and the relative contribution of green, yellow, and red wavelengths to total brightness (S1G, S1Y, and S1R respectively). The code for processing and analyzing spectral reflectance data is available at https://github.com/ksummers26/pavo-code.

#### Exome capture design

The large size and highly duplicated composition of the *R. imitator* genome (*12*) poses an obstacle to population genomic investigations into this species. We surmounted this hurdle by capturing and sequencing approximately 28 Mb of the *R. imitator* exome. This allowed us to obtain genomic-scale, short-read data for 124 individuals at coding regions of the genome potentially relevant to color phenotypes that we could reliably assemble and map. At the time of our study, there were no existing dendrobatid genomic resources to guide capture probe design, so we designed probes using *R. imitator* transcriptomes from different tissues and developmental stages.

In order to obtain transcriptomes for capture probe design, we first isolated mRNA from five tissues belonging to two young adult, Tarapoto-morph *R. imitator* (brain plus eyes, skin from the dorsal trunk, nape plus dorsal head, ventral jaw, and rear leg regions) as well as mRNA from entire tadpole skins sampled at weeks one (n = 1), two (n = 3), four (n = 3), and five (n = 3) of development (fig. S5). We prepared cDNA libraries with unique barcodes for each individual and tissue separately using Illumina TruSeq stranded total RNA kits. Various developmental stages were sequenced since we had no *a priori* knowledge regarding when color and pattern-related genes are expressed, though color becomes visible at around week five (fig. S5). Two library pools, one consisting of libraries from the young adults and another from all five developmental series, were each sequenced on two lanes of the Illumina HiSeq 2000 using 100 bp paired-end reads.

We assembled the sequencing reads from each cDNA library using Trinity (*63*), yielding 273,039 transcripts. Following the procedure of (*64*), we assigned Ensembl gene identifiers to 12,305 unique transcripts with a maximum BLASTX (*65*) e-value of 1e-10 and at least 50% sequence similarity to *Xenopus tropicalis* proteins downloaded from Ensembl (transcriptome annotation procedures are implemented in the script available at https://github.com/CGRL-QB3-UCBerkeley/MarkerDevelopmentPylogenomics/blob/master/5-Annotation). We also identified transcripts with unusually high differences in mapped read counts among the different young adult tissues and between the different developmental stages using EdgeR (*66*) that could suggest differential expression. Additionally, we considered any transcripts with TPM values, calculated with Kallisto (*67*), that were within the 95th percentile across all stages of development to be constitutively expressed genes potentially crucial for development. TransDecoder v 2.1.0 (*68*) was run on the unannotated transcripts with putative differential or constitutive expression patterns in order to extract those with open reading frames of at least 100 bp, from which we further trimmed away any non-gene-like portions. These trimmed, unannotated transcripts with interesting expression patterns along with all annotated transcripts were used as template sequences for capture probe design.

We applied a suite of filters to the set of transcripts used for probe design in order to optimize capture performance. This filtering entailed first performing a reciprocal blast of the transcripts and retaining one isoform per gene. For the retained transcripts, we trimmed each of the 3’ and 5’ untranslated regions (UTRs) to 500 bp in order to avoid designing probes on exceptionally long UTRs that could potentially represent assembly artifacts. We then masked out any known vertebrate repetitive elements and low-complexity regions from our target using repeatMasker v. 4.0.5 (*69*), since such elements could promote off-target capture. We retained only transcript regions consisting of at least 80 contiguous unmasked bases with GC content in the range of 40-70%, outside of which capture is ineffective (*70*). Lastly, since mitochondrial genes are unlikely to affect color phenotypes, we excluded all mitochondrial transcripts except for cytochrome B, which was kept to ensure some haploid representation for downstream quality control. Our final, filtered capture target included 13,265 genes spanning 28,281,490 bp. Among these genes, 10,904 were annotated with Ensembl IDs and accounted for 24,530,337 bp, while the remaining 2,361 unannotated genes with intriguing expression patterns totalled 3,751,153 bp in length. Custom capture probes were designed from this target sequence and synthesized as a NimbleGen SeqCap EZ Developer Library kit.

#### Genomic library preparation, exome capture, and sequencing

We extracted DNA from single toe clips for each *R. imitator* sample using either Qiagen DNeasy Blood and Tissue kits following the manufacturer’s protocols or a standard salt precipitation. For the salt precipitation method we first incubated each toe clip in 500 μL cell lysis buffer (1 mM Tris pH 8, 100 mM NaCl, 10 mM EDTA pH 8, 0.5% SDS) and 10 μL 20 mg/ mL proteinase K at 55 °C for 24 hours. After the tissues were completely digested we added 3 μL of 10 mg/mL RNase A to the samples and incubated them for an additional 30 minutes at 37 °C. Following this RNA digestion the samples were placed in a freezer at −20 °C for five minutes. We then added 200 μL of 5 M NaCl to each sample followed by vortexing for 20 seconds, and then placed them back in the freezer for another five minutes. Samples were then centrifuged at 11,000 RPM for eight minutes to pellet the proteins. The supernatant containing DNA was then transferred to new tubes, leaving behind the protein pellet, and 600 μL of 100% isopropanol was added to the tubes containing DNA, followed by 50 gentle inversions of the tubes. The samples were then placed in the freezer for one hour and then centrifuged at 10,000 RPM for 10 minutes to pellet the DNA. After pouring off the supernatant, we washed the DNA with 600 μL of fresh 70% ethanol. The tubes were inverted to dislodge the pellet in the 70% ethanol, and then centrifuged for eight minutes at 10,000 RPM. After centrifugation, the supernatant was poured off, leaving the DNA pellet behind, which was allowed to dry for at least 12 hours or until completely dry. Lastly, the samples were resuspended in 50 μL TE buffer.

For samples that had been extracted using the Qiagen kits, we performed an additional post-extraction RNase A digestion to ensure complete removal of RNA. Specifically, three μL cell L of 10 mg/mL RNase A was added to DNA suspended in Qiagen AE buffer and incubated at 37 °C for 30 minutes. The DNA was then recovered by adding 0.1x volume of 3M sodium acetate (pH 5.2) and two volumes of 100% isopropanol to the samples, followed by gently inverting the samples 50 times. The samples were then placed into a −20 °C freezer for 15 minutes. After removing the samples from the freezer, we centrifuged them at 10,000 RPM for 12 minutes at room temperature to pellet the DNA. We then discarded the supernatant and washed the DNA pellet by adding 600 μL cell L of fresh 70% ethanol to each sample followed by inverting the tubes to dislodge the pellet. The samples were then centrifuged at 10,000 RPM for 12 minutes. Lastly, we poured off the supernatant, which we allowed to dry for at least 12 hours. After the samples were completely dry, we eluted them in 50 μL cell L TE buffer.

We prepared genomic libraries for capture following the protocol of Meyer & Kircher (*71*). We started the library preparation with 0.92-1.4 μL cell g DNA per sample as measured by a Nanodrop fluorospectrometer, save for one sample (CH-14-5) for which we used 0.4 μL cell g DNA. The DNA for each sample was sheared using a Diagenode Bioruptor to an average fragment size of 250 bp for striped and banded samples and 300 bp for the admixed samples. For striped and banded samples this was achieved by shearing samples at the high Bioruptor setting for a total of seven minutes using cycles defined by 30 seconds of shearing followed by a 30-second pause (each sample was in the Bioruptor for a total of 13 minutes). The admixed samples were sheared at the medium Bioruptor setting for a total of four minutes using the same 30 seconds on / 30 seconds pause cycle scheme (each sample was in the Bioruptor for a total of eight minutes). Following each enzymatic reaction up until the indexing PCR step, all samples were cleaned using 1.6x volume Sera-Mag bead solution. A sample-specific, seven-nt barcode was incorporated into the library sequences for each sample via indexing PCR using 4 μL cell L of template library DNA and 10 PCR cycles under the reaction conditions specified in (*71*). We performed three independent indexing PCR reactions for the striped and banded individuals followed by bead purification using 1.3x volume Sera-Mag beads. For the admixed individuals we performed five independent PCR reactions, each of which were purified using 0.8x volume Sera-Mag beads. All of the PCR products for a respective individual were pooled across each of the independent PCR reactions.

Barcoded libraries for striped and banded samples were pooled equimolarly in batches of 22 individuals to generate three multiplexed pools for capture. All 58 admixed individual libraries were equimolarly pooled to form one multiplexed pool for capture. We isolated the targeted sequence from these capture pools using our custom *R. imitator* capture kit following NimbleGen’s protocols with slight modifications. Given the large genome size of *R. imitator*, we used approximately 2-3 times the amount of input DNA for hybridization compared to the 1 μL cell g called for by the NimbleGen protocol in order to preserve the complexity of the captured libraries: The three banded/striped captures used 2.522, 2.388, 2.663 μL cell g of DNA and the admixed capture used 3 μL cell g of DNA as measured by a Qubit fluorometer. We also doubled and tripled the amount of barcode blockers and COT-1 DNA used in the banded/striped and admixed hybridization reactions, respectively, relative to the amounts specified in the NimbleGen protocols. For COT-1 blocking DNA we used a cocktail of chicken Hybloc, human COT-1, and mouse COT-1 combined in equal amounts. All libraries were hybridized with the capture probes for 75 hours. The captured libraries were LM-PCR amplified in three separate reactions using 20 μL cell L of template DNA solution. The number of PCR cycles ranged from 11-14 cycles in order to yield DNA concentrations between 20-25 ng/μL cell L per reaction according to Nanodrop measurements. PCR products from the three reactions for each capture were pooled in equal amounts based on Qubit fluorometer measurements. Enrichment efficiency for the four capture experiments was assessed using qPCR with negative (off-target) and positive (targeted) control loci. The three banded/striped captures were pooled equally based on qPCR measurements for sequencing. The banded/striped multiplexed library and the admixed multiplexed library were sequenced on the Illumina HiSeq 4000 using 100bp and 150 bp paired-end reads, respectively.

#### De novo exome assembly

We applied quality controls to the sequencing reads using the program readCleaner (https://github.com/tplinderoth/ngsQC/tree/master/readCleaner) prior to assembly and mapping in the following order: 1) remove reads derived from PCR duplicates, 2) trim adapter sequences from the ends of reads, 3) trim low quality bases from the ends of reads in a two-step approach; first, trim bases with average Phred-scaled quality in 4 bp sliding windows below 20 from the ends of reads, and secondly, apply the BWA (*72*) algorithm implemented in cutadapt (*73*) using a minimum quality threshold of 20, 4) remove low complexity reads with a DUST score (*74*) above four, 5) merge read pairs that overlap by at least six base pairs and that have an observed expected alignment score (*75*) p-value less than 0.01, 6) remove reads that map to the human GRCh38 or *Escherichia coli* genome assemblies (NCBI GenBank accessions U00096, AE000111-AE000510), 7) remove reads shorter than 36 bp and/or have at least 50% of their sequence comprised of Ns.

We assembled quality-controlled reads from six *R. imitator* individuals (three banded morph: ET13012, ET13017, ET13018, and three striped morph: ET13029, ET13032, ET13033) separately with SPAdes v 3.10.0 (*76*) under default parameters. Contigs with at least 80% sequence identity to targeted *R. imitator* transcripts, determined using blastn, were clustered using CD-HIT (*77, 78*) and merged with CAP3 (*79*). All resulting contigs belonging to the same gene were concatenated together in syntenic order with interspersed runs of 39 Ns to prevent reads from mapping across the gaps. Scripts used for assembly are available at https://github.com/CGRL-QB3-UCBerkeley/seqCapture.

#### Variant discovery

We mapped *R. imitator* reads with at least 30 high-quality bases (NovoAlign option -l) to the *de novo* exome assembly using NovoAlign v 3.04.06 (*80*), with the highest acceptable alignment score for the best alignment set to 150. Only reads with a minimum Phred-scaled mapping quality of 20 were retained. The NovoAlign average insert size and standard deviation mapping parameters were set according to values determined from bioanalyzer traces for the captured genomic libraries. Banded and striped individuals were mapped using an average insert size of 220 bp and a standard deviation of 58, while for admixed individuals the insert size was set to 271 bp with a standard deviation of 63. We used SAMtools v 1.11 (*81*) to merge unpaired and paired alignments from the same individual. We mapped RNAseq reads from the model species, *R. summersi* and *R. variabilis,* to the *R. imitator* exome assembly using *bwa mem* v. 0.7.17 (*82*) with default parameter settings. For the models, we used SAMtools view to retain only primary alignments for which the reads mapped in proper pairs while excluding all reads that were PCR or optical duplicates or which failed quality checks (-F 1804).

Prior to identifying variants, we produced quality control masks denoting genomic regions where variant calling would be reliable. This was achieved by first generating an all-sites (variable and monomorphic sites) VCF containing all *R. imitator* and model species individuals, which included coverage and map quality information at every site, as well as p-values for tests of various biases between reference and alternate alleles called using bcftools v 1.10.2 (*83*). Given the high levels of duplication throughout the *R. imitator* genome we were particularly cautious to mask regions refractory to short reading mapping. We performed this masking using ngsParalog (https://github.com/tplinderoth/ngsParalog), which was ran for the banded, striped, and admixed *R. imitator* populations separately. Specifically, the ngsParalog calcLR function calculates a likelihood ratio (LR) statistic for testing whether within-individual read proportions and sample-wide genotype likelihoods at a site are consistent with reads sourced from multiple loci. This enabled identification of chimeric regions in the exome assembly. We annotated the all-sites VCF with these likelihood ratios, and used bcftools to mask out multiallelic sites and indels, as well as sites with an RMS mapping quality score of less than 30, total site coverage greater than the 99.8 exome-wide percentile, p-values for either strand, base quality, map quality, or end-distance bias that were less than 1e-100, p-values for excess heterozygosity from called genotypes less than 1e-3, and ngsParalog LRs greater than 500 (approximately the 98th genome-wide percentile) in any of the *R. imitator* populations.

In addition to the above filters we required at least 15 individuals from each of the striped, banded, and admixed *R. imitator* populations to be covered by at least three reads at a site in order for the site to pass masking. We also imposed three levels of coverage requirements for the model species according to their relevance to particular analyses in order to maximize the amount of quality data while minimizing bias (fig. S6): 1) no coverage requirement for GWAS, which were performed among *R. imitator* only, 2) one individual of either model species with data for comparative population genetic analyses (e.g. PCA) and tests for introgression and selection, 3) all individuals from both model species with data for phylogenetic inference. We called SNPs based on a maximum p-value of 1e-6 for the likelihood ratio test of the minor allele frequency being greater than zero among all *R. imitator* and model species at unmasked sites using ANGSD. Estimation of low frequency variants was particularly sensitive to missing data (fig S7), therefore we masked variable sites with minor allele frequency less than 2% in *R. imitator* (five minor alleles) from all analyses. The distribution of allele frequencies estimated with ANGSD was robust to missing data at frequencies above this cutoff.

#### Population genetic characterization

We estimated levels of genetic diversity within *R. imitator* populations in terms of the number of segregating sites, *S*, Watterson’s estimator, *θ*_Watterson_, and nucleotide diversity, *π*, in ANGSD based on the genotype-likelihoods of individuals in our sample. The latter two diversity estimates were estimated using an empirical Bayes procedure that uses each population’s respective exome-wide, folded site frequency spectrum (SFS, also calculated with ANGSD) as a prior (fig. S6).

Genomic relationships among *Ranitomeya* individuals were assessed with principal components analysis (PCA). We first calculated the genetic covariance matrix among all individuals with ngsCovar (*85*) based on individual’s genotype posterior probabilities under a Hardy-Weinberg allele frequency prior in ANGSD. We decomposed this matrix using the eigen() function in R v. 3.6.3 to obtain the eigenvectors and their corresponding eigenvalues for the PCA. We also repeated this analysis on *R. imitator* only. We quantified pairwise genetic divergence between the different *R. imitator* populations using Reynold’s F_ST_ estimator (*86*) with ANGSD. As a prior on the distribution of joint allele frequencies for calculating F_ST_, we used the unfolded joint SFS between the respective pair of populations considering the reference base as ancestral (mispolarization with respect to ancestral and derived allelic state does not matter for this calculation). Lastly, we estimated the proportions of striped and banded *R. imitator* ancestry of individuals sampled from the introgression zone using NGSadmix (*57*). We ran NGSadmix on genotype likelihoods from ANGSD for all striped, banded, and admixed *R. imitator* individuals, specifying 2-5 ancestral populations (K parameter) (fig. S8).

#### Divergence mapping

We expect to see increased divergence between the striped and banded morphs of *R. imitator* at loci that contribute to the color and pattern differences between them. However, we expect the two morphs to be somewhat differentiated even at random loci. In order to identify loci that are more differentiated than expected given the genome-wide distribution of allele frequency differences, we used the test statistic *δ* =|*p̂*_1_ − *p̂*_2_|, where *p̂*_1_ and *p̂*_2_ are the estimated allele frequencies in the banded and striped population, respectively. To assign p-values we simulated data of the joint allele frequencies in the two populations using the Balding-Nichols model (*15, 16*). Given an ancestral allele frequency, *p*_0_, the allele frequencies under the Balding-Nichols model in two derived populations with genome-wide divergence expressed in terms of F_ST_, are two independent draws from the beta distribution β(*p*_0_(1-F_ST_), (1-*p_0_*)(1-F_ST_)/F_ST_)). Using two draws from this distribution, (p_1_,p_2_), we modeled allele sampling based on these true frequencies with a binomial distribution: *p̂*_1_ ∼ B(2*n*_1_,*p*_1_) and *p̂*_2_ ∼ B(2*n*_2_,*p*_2_), where *n*_1_ and *n*_2_ are the diploid sample sizes from populations 1 and 2 respectively. The approach for approximating p-values for observed allele frequency differences, *δ*_obs_, based on these expected allele frequency differences under the Balding-Nichols model is outlined below:

1) We simulated distributions of *δ* under the Balding-Nichols model in bins of different ancestral allele frequencies (*p*_0_). We used bin widths for *p*_0_ equal to 1/(2(*n*_1_+*n*_2_))= 0.007575758 for our sample sizes. Each distribution of *δ* was generated by sampling *p*_0_ uniformly in the range of its corresponding bin one million times. Note that we only need to consider allele frequencies up to 0.5 because distributions of *δ* are symmetric for higher frequencies, resulting in a total of *n*_1_+*n*_2_ bins. We also discretized the distributions of *δ* into *k* bins of width argmax(1/2*n*_1_, 1/2*n*_2_) = 0.01515152 for our sample sizes and defined *P* (*δ_r_* | *p*_0_ *bin i*) as the proportion of *δ* values in the simulations with values of *p*_0_ in *p*_0_-bin *i* that fell in *δ* -bin *r, r ∈*{1,… *, k* }.

2) We approximated the distribution of *p*_0_, i.e. the probability of *p*_0_ falling in the *i*th bin, empirically as

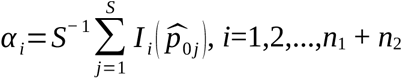

where *I_i_* (*p̂*_0_*_j_*) is an indicator of *p̂*_0_*_j_* falling in the *i*th bin, *p̂*_0_*_j_* is the value of (*n*_1_*p̂*_1_ + *n*_2_*p̂*_2_)/(*n*_1_ + *n*_2_) calculated for the *j*th SNP, and *S* is the total number of sites.

3) Denote an observed allele frequency difference falling in *δ*-bin *j* by *δ*_obs,j_. We then approximated the p-values as

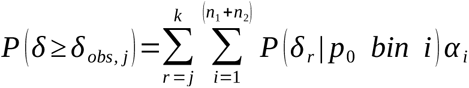

While this simple model may not adequately capture all of the intricacies of a more parametric demographic model, it provides a convenient statistical framework for identifying divergent loci. This procedure for approximating p-values is implemented in https://github.com/tplinderoth/PopGenomicsTools/blob/master/betaAFOutlier.R.

Our estimated genome-wide F_ST_ in the Balding-Nichols model was obtained using ANGSD. Observed allele frequency differences between the banded and striped morphs were calculated using allele frequencies estimated for each of the respective populations with ANGSD -doMaf 1 and setting the major allele to the reference allele with -doMajorMinor 4. P-values were approximated for the observed allele frequency differences using the procedure outlined above, which we used to test for significant *δ* s at a 0.05 level after a Bonferroni-adjustment for the number of tested SNPs.

#### Association mapping in the admixed population

As a follow-up on the candidate genes identified from the divergence mapping, we performed a second, independent genome-wide scan among admixed individuals of *R. imitator*. We used a general linear modeling (GLM) approach to quantify the effect of genotypes on continuously measured phenotypes while accounting for covariates (*87*). For each SNP, we estimated the genotypic effect on pattern orientation (PT_rot_), dorsal body color (DC_a_), and hindlimb color (LC_a_) (see “Phenotyping” of Materials and Methods) based on the following models with *N* denoting a normal distribution with variance *σ*^2^

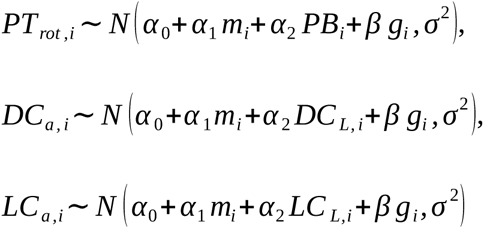

The genotype of individual *i*, *g_i_ ∈*{0,1,2}, can have 0, 1, or 2 copies of the minor allele. The admixture proportion, *m_i_*, is the proportion of banded ancestry for individual *i* estimated with NGSadmix, and is equal to 1 – striped ancestry proportion. This covariate controls for confounded associations from differences in genome-wide levels of striped and banded ancestry. DC_L_ and LC_L_ are lightness values quantified from digital images in CIELAB color space to control for lighting effects. The GLMs for color quantified along the red-green CEILAB axis (a*) included lightness as a covariate in order to account for non-standardized lighting conditions across frog photos. We compared twice the log likelihood ratio of a genetic association, which is the ratio of the log likelihood at the maximum likelihood estimate of *β* and the likelihood when *β*=0, to a *X*^2^_1_-distribution to obtain p-values for each SNP.

We combined the p-values for each SNP from the color-morph divergence test, *p_d_*, with the p-values from the GLM-based tests of association in the admixed population, *p_a_*, using Fisher’s method (*24*). This test statistic is defined as *X*^2^_*i*_ = −2 [log(*p_d,i_*)+log(*p_a,i_*)] for each SNP, *i*, in the set of loci common to both tests. While the distribution for these combined p-value statistics, X^2^, would theoretically follow a *X*^2^_4_-distribution under the null when each independent test aligns with the null (no association), we observed subtle genome-wide inflation (fig. S9). Consequently, we fit the degrees of freedom, *k ∈* ℝ^+^, of a *X*^2^*_k_*-distribution to our observed X^2^ distribution in R v. 3.6.3. and used this fitted distribution as our null for calculating p-values (fig. S9). This assumes that nearly all SNPs in the genome are not associated with the tested phenotype and is otherwise conservative. We tested for significant X^2^ values at the 0.05 level for falsely rejecting the null of no genetic associations after correcting for the number of tested SNPs using a Bonferroni adjustment.

#### Phenotypic variance explained by candidate SNPs

We estimated how much phenotypic variation in pattern orientation (*PT_rot_*), dorsal body red-green color (*DC_a_*), or hindlimb red-green color (*LC_a_*) among *R. imitator* from the admixture zone was explained by genotypes at the most highly associated color gene SNPs using linear models in R v. 3.6.3. Specifically, the percentage of variance explained for each of the phenotypes was quantified as 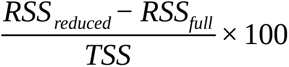, where *TSS* stands for the total sum of squares and *RSS_full_* and *RSS_reduced_* denote the residual sum of squares for the linear models with and without a candidate SNP predictor variable, respectively. In the full models, for each candidate SNP, the phenotypes of individual *i* were modeled as

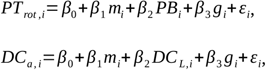

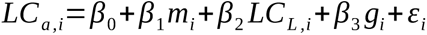

where the genome-wide admixture proportion, *m_i_*, is individual *i*’s banded ancestry proportion (equal to 1 – striped ancestry proportion) and *g_i_ ∈*{0,1,2} is their number of minor alleles at the candidate SNP. *PB*, *DC_L_*, and *LC_L_* are covariates for the black proportion of the body dorsum and the CIELAB lightness values for the body dorsum and legs, respectively, as defined in the “Phenotyping” section of the Methods. *β*_0_ is an intercept term, *β*_1,..,3_ are effect sizes of the predictor variables, and *ε* is the model error. The reduced models were exactly the same, but lacked the candidate SNP predictor, *G*. The genotypes at candidate SNPs were called using ANGSD -doPost 1, -postCutoff 0.95, and -doGeno, which calls a genotype if its posterior probability is above 0.95, otherwise it is considered missing. The *β* parameters of the linear models were estimated using the lm() function and the incremental variance explained by inclusion of *G* was assessed using anova() to compare the full and reduced models in R.

#### Linkage analyses in lab-reared pedigrees

We genotyped candidate coloration gene SNPs identified from the genome-wide scans (or physically proximal SNPs in the same genes that were strongly linked to candidate SNPs; whichever we could design the most reliable primers for) in the captive pedigree of *R. imitator* in order to conduct linkage analyses. We did not conduct analyses on *krt8.2* in the pedigree due to technical difficulties sequencing this locus stemming from limited sequence for primer design around the candidate SNP of interest. We amplified target genic regions in 10 µL PCR reactions consisting of final concentrations of 1 µL DNA template, 1X Invitrogen *Taq* DNA polymerase PCR buffer, 2.5 mM MgCl_2_, 0.05 mM of each dNTP, 0.2 µM of each of the forward and reverse primers (table S12), and 0.5 µL *Taq* DNA polymerase. Thermal conditions for PCR amplification consisted of an initial denaturation for 2 minutes at 94°C, followed by 30 cycles of, followed by 30 cycles of 94°C, followed by 30 cycles of for 30 seconds, 56°C, followed by 30 cycles of for 40 seconds, and 72°C, followed by 30 cycles of for one minute, and then lastly, a final extension at 72°C, followed by 30 cycles of for 10 minutes. After DNA amplification, remnant primers and dNTPs were hydrolyzed using ExoSAP-IT by incubating at 37°C, followed by 30 cycles of for 30 minutes, followed by 80°C, followed by 30 cycles of for 15 minutes in a thermalcycler. Amplicons were cycle-sequenced in 10 µL reactions consisting of 2 µL of purified PCR product and final concentrations of 0.25X BigDye Terminator 3.1 ready reaction mix, 0.75X BigDye Terminator v3.1 sequencing buffer, and 0.3 µM primer. Sequencing was carried out using an initial denaturation at 96°C, and 30 cycles of 96°C 5 seconds, and 60°C, followed by 30 cycles of for four minutes. Cycle sequencing products were strained through a sephadex matrix using centrifugation at 3000 RPMs for two minutes, and then the purified products were sequenced using an Applied Biosystems 3730 Genetic Analyzer.

Sequence chromatograms were analyzed in Geneious v. 4.8.5 (*88*). We first manually inspected and trimmed low quality positions with ambiguous peaks from the ends of both the forward and reverse sequences. Any sequences containing ambiguous peaks throughout their entire lengths were discarded. The forward and reverse sequences of each sample were then aligned and manually checked for genotyping errors, ensuring correct calls for both homozygous and heterozygous sites by retaining only genotype calls where both the forward and reverse sequence matched and were high quality with clear peaks. Sites were translated into exomic coordinates by aligning the forward-reverse consensus sequences for each sample to the respective candidate gene contigs of the exome assembly.

Linkage analyses were performed using the Merlin software (*89*). We used the default settings, aside from changing the complexity threshold to allow more complex (larger) families. We tested for a genetic association with pattern orientation at candidate SNPs in *mc1r, asip,* and *bsn*. Some phenotypic measures of color were correlated (fig. S10), and so we used a hierarchical testing scheme to minimize redundant testing. We first tested for an association between candidate SNPs in *mc1r, asip, bsn*, and *retsat* with dorsal body and hindlimb brightness. If we failed to find a statistically significant association, then we performed subsequent tests of chroma in order to rigorously test for any association with color. Association analyses were performed separately for each body region and candidate gene SNP. Logarithm of the odds (LOD) scores were calculated, with positive scores indicating that allele sharing among individuals with similar phenotypes is higher than expected in the absence of a true association between genotype and phenotype. We controlled for falsely identified associations from multiple testing using the procedure of Benjamini-Hochberg (*90*), specifying a false discovery rate threshold of 10%. Though this procedure assumes independent tests and some of the tested phenotypes were correlated (fig. S1), this approach for controlling for false positives is considered to be mostly robust to non-independence (*91*).

#### Phylogenetic relationships and tests for introgression

We estimated neighbor-joining trees to visualize the evolutionary relationships between *R. imitator* and the model species across the genome and at candidate color genes in order to discern whether *R. imitator* acquired color alleles from the model species. Specifically, we calculated a pairwise genetic distance matrix among all individuals based on their genotype likelihoods using ngsDist (*92*) set to all defaults. For each SNP, the likelihoods of all nine possible genotypes for a pair of individuals were considered. We performed bootstrap sampling of single sites to generate 100 bootstrap replicates using ngsDist for node support. The resulting distance matrix served as input to construct a phylogenetic tree with FastME (*93*) using the BIONJ algorithm and SPR tree topology improvement. The tree was visualized in R using the ‘phytools’ package (*94*).

We used ANGSD -doAbbababa2 (*95*) to test for introgression from the model species into *R. imitator*. Analyses were restricted to reads from input BAM files with minimum base quality of 20 and which aligned uniquely with a map quality of at least 20. We ran -doabbbabba2 using all of the reads that passed quality control at each site (*-*sample 0) up to total depth of 7,100 (-maxDepth 7100) and the default block size (-blocksize 5 Mb) which effectively counts ABBA and BABA patterns for entire contigs (genes in our case) if they are less than 5 Mb long. For the ABBA-BABA test, we specified banded *R. imitator,* striped *R. imitator*, and *R. variabilis* as the H1, H2, and H3 ingroups and *R. summersi* as the outgroup. Given this configuration, excess ABBA patterns would result in a positive D-statistic, which, if significantly greater than zero, would suggest introgression between the mimics and models. We used the estAvgError.R script packaged with ANGSD to calculate the average exome-wide D-statistic and to test for whether it was significant at the 0.05 level for falsely rejecting the null hypothesis of no admixture.

#### Selection

We used Hudson–Kreitman–Aguadé (HKA) tests (*28*) to determine whether the levels of polymorphism within *R. imitator* compared to substitutions from the model species were consistent with neutral evolution at exons containing SNPs associated with coloration. These tests were performed for three different sets of ingroups: the banded *R. imitator* morph, the striped *R. imitator* morph, and the banded and striped *R. imitator* morphs combined. The outgroup for this test consisted of the model species *R. variabilis* and *R. summersi* together. We used ANGSD to identify the minor allele from the genotype likelihoods (-GL 1 and - doMajorMinor 1) for all striped and banded *R. imitator*, *R. variabilis*, and *R. summersi* jointly (hereon called the “joint minor allele”) at all high-quality sites across the genome where there was data for at least 15 individuals from each of the striped and banded morphs and one individual from either model species (see “Variant Discovery” of the Materials and Methods). We called a site variable if the likelihood ratio test p-value (outputted with -SNP_pval) for whether the joint minor allele frequency (estimated with -doMaf 1) was different from zero was at or below 0.01. These SNPs were considered polymorphic within a *R. imitator* morph if the morph-specific minor allele frequency (MAF) was at least 1% and polymorphic for the striped and banded morphs combined if the site was polymorphic within at least one of the morphs or a fixed difference between them. Similarly, a SNP was considered variable in one of the model species if the species-specific MAF was at least 1% and a SNP was variable among the model species (the outgroup for all HKA tests) if the site was variable within at least one of the model species or a fixed difference between them. The HKA procedure used the subset of SNPs that were either polymorphic in the ingroup or substitutions from the outgroup and omitted any sites that were polymorphic in the outgroup. All within-morph and within-species MAFs were estimated using ANGSD -doMaf 1. All genotype likelihood and allele frequency calculations in ANGSD were limited to bases with minimum quality scores of 20 and reads that mapped in a proper pair with a minimum map quality of 20. Minor allele frequency estimate cutoffs needed to be used to identify variable sites since hard calling genotypes would diminish accuracy in the context of our medium-depth sequencing data.

We calculated the HKA test statistic, *X*^2^, as in (*96*). Specifically, for each set of tests involving a different ingroup we used the fraction of SNPs across the genome that were segregating in the ingroup, and substitutions from the outgroup, as estimates for the neutral rates of ingroup polymorphism, *p*_s_, and substitutions, *p*_d_, respectively. Since we condition on sites that are either segregating in the ingroup or substitutions from the outgroup, *p*_d_ = 1 - *p*_s_. For a locus consisting of *L* conditional SNPs, the expected number of segregating sites was calculated as *e*_s_ = *Lp*_s_ and the expected number of substitutions as *e*_d_ = *Lp*_d_. The HKA test statistic for a particular locus was calculated as 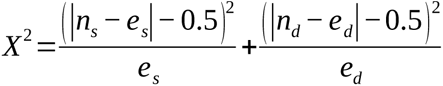, where *n*_s_ and *n*_d_ are the observed number of ingroup segregating sites and substitutions from the outgroup at the locus in question, respectively. Subtracting 0.5 from the differences of the observed and expected counts imposes the Yate’s correction for continuity such that the distribution of the *X*^2^ statistic can be better approximated by a *χ*^2^ distribution when at least some counts are small (*97*). We tested for neutral evolution at the exons (and whatever flanking sequence that was captured with them) containing phenotype-associated SNPs of interest.

Factors such as linkage-disequilibrium among SNPs and complex demographic histories can cause the null distribution for the *X*^2^ statistic to deviate from a theoretical *X*^2^_1_ distribution. To account for this we used the qq.chisq() function of the snpStat R package (*98*) to calculate a *X*^2^ correction factor, *λ*, based on the genome-wide distribution of *X*^2^ statistics, which comprised all *X*^2^ statistics for every exon in our exome assembly. *λ* was calculated as the ratio of the means between the observed genome-wide *X*^2^ and expected *X*^2^ distributions after trimming values above the median as per the qq.chisq “trim” argument default. Values of *λ* were, respectively, 0.46, 0.60, and 0.56 for the tests using the banded morph ingroup, striped morph ingroup, and collective banded and striped morph ingroup. After scaling the candidate exon *X*^2^ statistics by the respective values of *λ*, we compared them to a *X*^2^_1_ distribution in order to obtain p-values, which we adjusted based on the number of tested exons for a given ingroup using the procedure of (*90*).

We also tested for neutrality at the candidate exons containing phenotype-associated SNPs using the program HKA (*99*). Given *m* loci, the HKA statistic in this case was calculated as 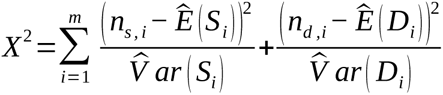, where *n*_S,i_ and *n*_d,i_ are the observed number of ingroup segregating sites and substitutions from the outgroup, respectively, at locus *i*. *Ê*(*S_i_*) and *V̂ ar*(*S_i_*) are estimates of the expectation and variance for the number of ingroup segregating sites at locus *i*, and *Ê*(*D_i_*) and *V̂ ar* (*D_i_*) are the expectation and variance for the number of substitutions between the ingroup and outgroup at locus *i.* These expectations and variances are obtained using estimates of the split time between the ingroup and outgroup, *T*, and the population-scaled mutation rate, *θ_i_*, for each locus *i*, under a standard neutral Wright-Fisher infinite sites model assuming equal population sizes for both the ingroup, outgroup, and ancestral population.

We calculated the HKA *X*^2^ statistic using coalescent simulations for the set of candidate gene exons containing phenotype-associated SNPs for ingroup configurations consisting of the banded and striped *R. imitator* morphs separately, and for the two morphs together. For each set of tests (ingroups), a null distribution for the *X*^2^ statistic was obtained using 1000 neutral coalescent simulations under the assumptions of the null HKA model (*28*) and estimates for *T* and *θ_i_*. P-values were calculated as the proportion of the respective null simulations greater or equal to the observed *X*^2^ statistic.

### Supplementary Text

#### Exome assembly and variant discovery

In order to map loci underlying color traits in the complex, 6.8 Gb (*12*) genome of *Ranitomeya imitator,* we designed custom sequence capture probes for 13,265 genes spanning 28.28 Mb of the exome from a developmental series of *R. imitator* transcriptomes. We used this capture system to obtain moderate-depth sequencing data (mean 12x) for 124 *R. imitator* individuals (33 banded, 33 striped, 58 admixed; Fig. 1). Sequencing data from three banded and three striped individuals were used to assemble 76.4 Mb of the *R. imitator* exome. This assembly, with an N50 of 786 bp, encompasses 13,084 (98.6%) of our targeted genes. The average contig length per gene was 5,825 bp. We mapped the sequence capture reads from all *R. imitator* samples and RNAseq reads from the model species, *R. summersi* and *R. variabilis*, to this exome assembly and jointly called 928,822 high-quality, transspecies SNPs after discarding sites segregating with a MAF below 2% in *R. imitator*. The conditional removal of sites that segregate within *R. imitator* at low frequencies was due to the fact that estimates of allele frequencies below 2% in *R. imitator* were sensitive to differences in quality control thresholds (fig. S7). Among *Ranitomeya*-wide SNPs, 295,279 were polymorphic in *R. imitator*.

#### Population genetic characterization of *Ranitomeya imitator*

Greater spread among striped compared to banded *R. imitator* individuals across the first two principal components of the PCA based on *R. imitator* genome-wide variation (Fig. 2A, right panel) reflects higher genetic diversity in the striped population: nucleotide diversity, *π* = 1.56*×*10^-3^, *θ*_Watterson_ = 1.38*×*10^-3^, 20,751 private SNPs among 33 individuals versus banded *π* = 9.56*×*10^-4^, *θ*_Watterson_ = 9.34*×*10^-4^, 8,835 private SNPs among 33 individuals. The genetic diversity of the population in the admixture zone (*π* = 1.31*×*10^-3^, *θ*_Watterson_ = 1.30*×*10^-3^, 25,054 private SNPs among 58 individuals) is slightly lower than that of the striped. Individuals from this population are positioned between the striped and banded populations along PC1, consistent with their geographic location and mixed ancestry from these pure-morph populations (*10*). This agrees with estimates of ancestry proportions based on two ancestral lineages (Fig. 2B).

#### Phenotypic variance explained by candidate SNPs

We quantified how much phenotypic variance in the wild, admixed population was explained by genotypes at the candidate SNPs with the largest *X*^2^ (Fisher’s combined test statistic for the divergence and admixture zone mapping experiments) for each respective phenotype, after accounting for differences in genome-wide ancestry among individuals and technical covariates (table S8). Candidate SNPs in the pattern-associated genes *bsn*, *mc1r*, and *asip* explained 9%, 9%, and 4% of the variation in pattern orientation, respectively. These genes were also associated with body color, with the highest *X*^2^-ranked SNPs in *bsn*, *mc1r,* and *asip* explaining 7%, 3%, and 2% of the color variation along the CIELAB (*57*) red-green axis, while the other body color candidate gene, *retsat*, explained 12%. The highest *X*^2^-ranked hindlimb color SNPs in *krt8.2, mc1r,* and *asip* explained 18%, 8%, and 1% of the hindlimb red-green axis variation.

#### Linkage analysis in a captive *R. imitator* pedigree

Each of the genotyped SNPs from all of the pattern candidate genes, *mc1r*, *asip*, and *bsn*, showed significant associations with pattern orientation among pedigree frogs (table S9, LOD range 0.93-2.54, q-value range 0.003 - 0.091).

The dorsal body colors in the pedigree frogs ranged from yellow to reddish/orange (Fig. 3A). Brightness (B2) and yellow wavelength chroma (S1Y, 550-625 nm) measured from the body dorsum were uncorrelated (Pearson’s r = 0.18, t = 1.6, two-tailed p-value = 0.1, see also fig. S10), so we tested for associations between both of these color features and SNPs in candidate body color genes (*mc1r, asip, retsat, bsn*). *Asip* was significantly associated with brightness (LOD = 2.47, q-value = 0.003) but not yellow intensity (LOD = 0.17, q-value = 0.432), whereas *retsat* showed a highly significant association with dorsal yellow intensity (LOD = 3.49, q-value = 8.54×10^-4^), but no association with brightness (LOD = 0.33, q-value = 0.342). Neither *mc1r* or *bsn* were significantly associated with brightness or yellow intensity in the pedigree (LOD scores ranging 0.09-0.48, q-values = 0.238-0.571).

There is pronounced variation in hues on the hindlimbs of the pedigree frogs (Fig. 3A), with colors ranging from yellow/orange (as in the wild banded population) to bluish/greens as in the striped population (Fig. 1). Accordingly, we measured both mean brightness (B2) and green wavelength intensities (S1G, 400-510 nm) from the hindlimbs. The green intensities are significantly correlated with brightness (Pearson’s r = 0.47, t = 4.2, two-tailed p-value = 7.6×10^-5^, fig. S10), so we only tested for associations with green wavelength if no significant associations with brightness were detected, to minimize redundant testing. We tested SNPs in the hindlimb color candidate genes *mc1r* and *asip* and found that the *mc1r* SNP was significantly associated with brightness (LOD = 1.13, q-value = 0.064). While *asip* was marginally associated with brightness (LOD = 0.62, q-value = 0.182), it showed no evidence of affecting green wavelengths (LOD = 0.0, q-value = 0.950). We did not test for associations between *krt8.2* and hindlimb color in the pedigree due to technical difficulties designing reliable sequencing primers based on the restrictive amount of assembled sequence flanking the candidate *krt8.2* SNP.

#### Tests of neutral evolution at candidate color genes

We used the Hudson–Kreitman–Aguadé (HKA) test (*28*) to determine whether levels of polymorphism within *R. imitator* relative to divergence from the model species was consistent with neutral evolution at the candidate gene exons that contained the strongest phenotype-associated SNPs (table S11). These tests rejected neutrality for the second exons of *mc1r* (*X*^2^*_imitator_* = 6.25, q = 0.032) and *bsn* (*X*^2^*_imitator_* = 18.35, q = 1.101e-4) and exon 11 of *retsat* (*X*^2^*_imitator_* = 5.82, q = 0.032). Furthermore, we found evidence that some of these exons evolved non-neutrally within the different *R. imitator* morphs (table S11). For the banded morph, exons containing associated SNPs showed significant departure from neutrality for all candidate genes, except *krt8.2*: *mc1r* exon 2 (*X*^2^*_banded_* = 22.44, q = 1.304e-5), *asip* exon 2 (*X*^2^*_banded_* = 4.67, q = 0.046), *bsn* exon 2 (*X*^2^*_banded_* = 14.28, q = 4.717e-4), *retsat* exon 11 (*X*^2^*_banded_* = 9.94, q = 0.003). For the striped morph we found significant departure from neutral patterns at *bsn* exon 2 (*X*^2^*_striped_* = 15.34, q = 5.394e-4), while *mc1r* exon 2 (*X*^2^*_striped_* = 4.46, q = 0.069) and retstat exon 11 (*X*^2^*_striped_* = 4.57, q = 0.069) approached the 5% false discovery significance level. We note that, in all cases where exons significantly deviated from neutral expectations, we observed decreased polymorphism within species or populations compatible with selective sweeps (table S11).

In these analyses we used genomic control to adjust for the fact that SNPs in linked sites are correlated and that the nominal p-values using standard tests of homogeneity, therefore, will not be calibrated accurately (*28*). However, we also used an alternative approach for obtaining p-values based on coalescent simulations under the standard neutral coalescent model. In close agreement with the HKA results based on *X*^2^_1_ approximations, there was significant evidence of non-neutral evolution at candidate gene exons containing associated SNPs (*mc1r* exon 2, *asip* exon 1, *asip* exon 2, *bsn* exon 2, *retsat* exon 11, *krt8.2* exon 3) in the test involving both morphs combined (*X*^2^*_imitator_* = 10.78, p = 0.03) and in the banded morph (*X*^2^*_banded_* = 18.70, p = 0.01) while neutral evolution was nearly rejected for the striped morph (*X*^2^*_striped_* = 9.42, p = 0.06).

**Fig. S1.**
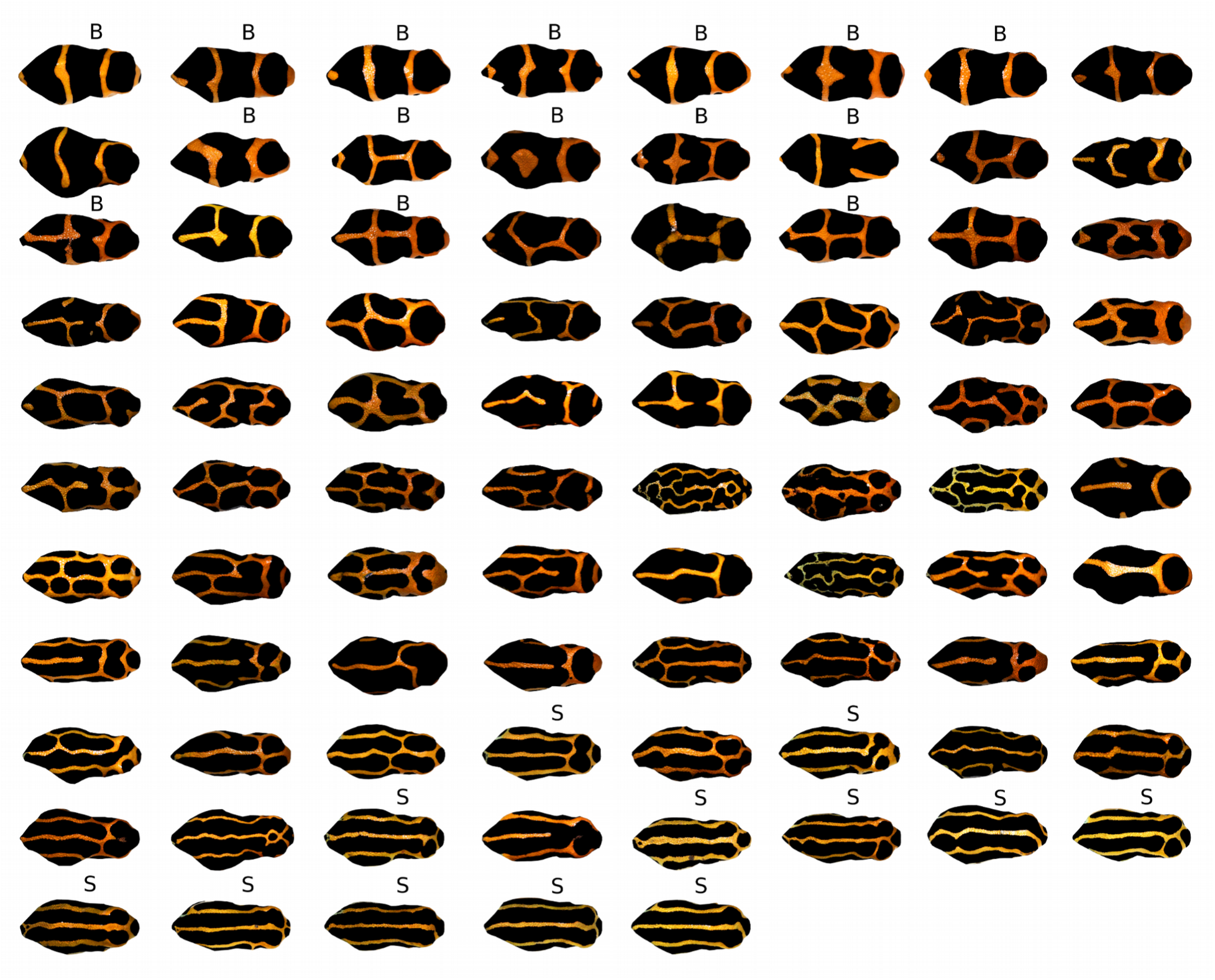
Images of *Ranitomeya imitator* sorted according to pattern orientation score. Digital images from photographs of the dorsal surface of *R. imitator* are sorted according to the pattern orientation score, with the score increasing from the top to bottom row as well as left to right; the top-left frog has the lowest (most banded) score while the bottom-right frog has the highest (most striped) score. Frogs labeled with a “B” and “S” are from the banded and striped morph populations respectively, and all other frogs were sampled from the admixture zone.

**Fig. S2.**
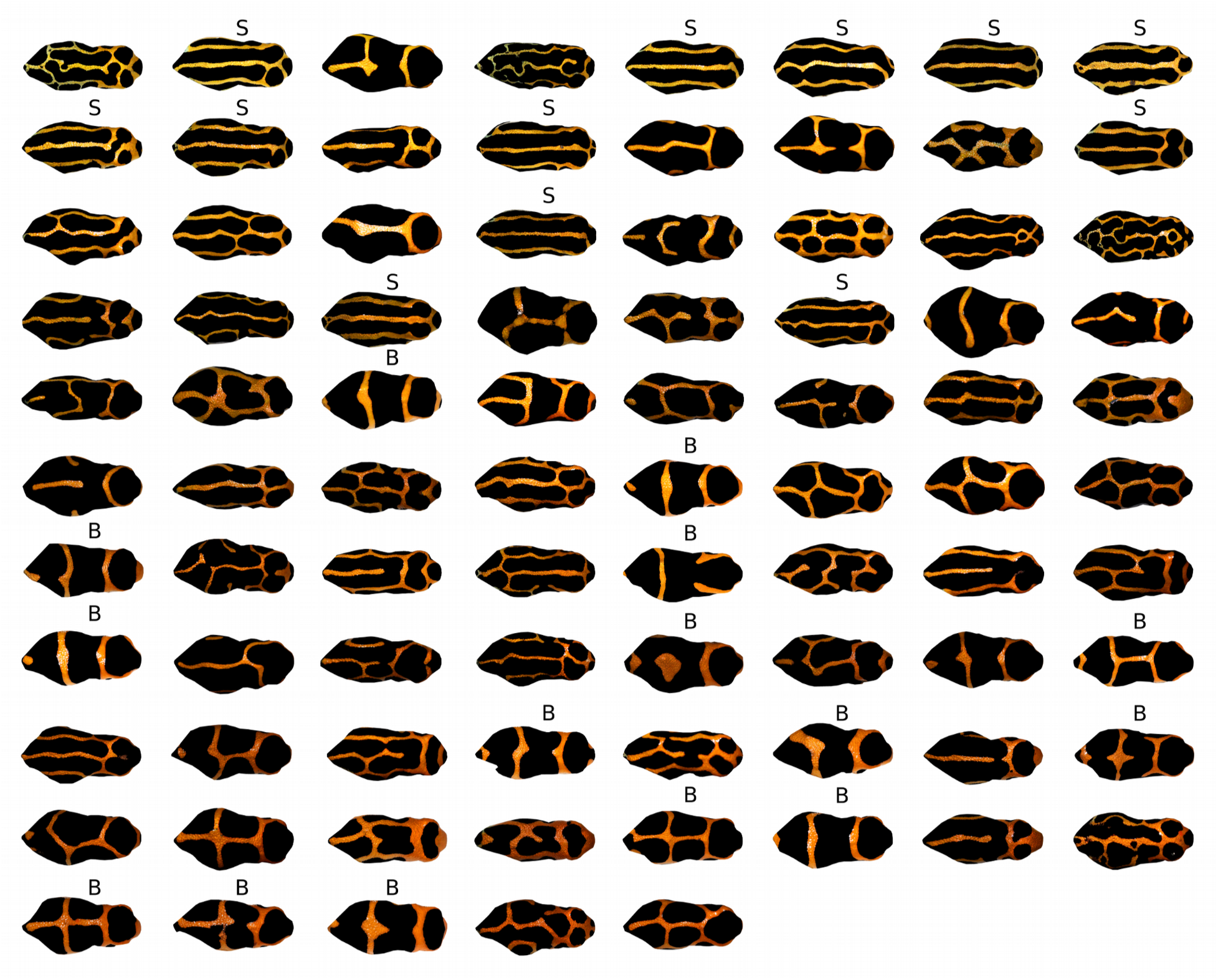
Images of *Ranitomeya imitator* sorted according to body dorsum a* color value. Digital images from photographs of the dorsal surface of *R. imitator* are sorted according to their CIELAB a* (green-red axis) value, with a* increasing from the top to bottom row as well as left to right; the top-left frog has the lowest (most green) value while the bottom-right frog has the highest (most red) value. Frogs labeled with a “B” and “S” are from the banded and striped morph populations respectively, and all other frogs were sampled from the admixture zone.

**Fig. S3.**
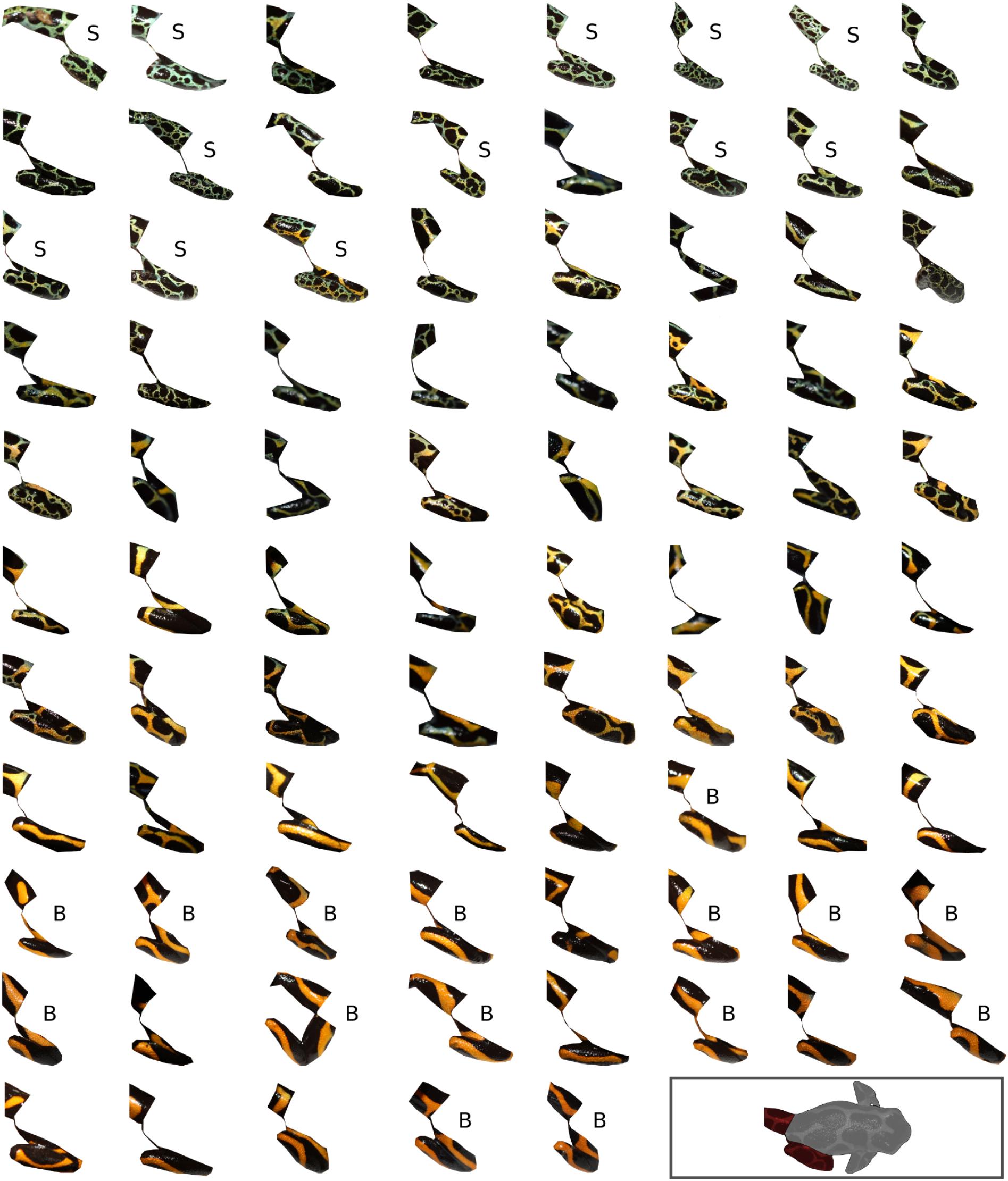
Images of *Ranitomeya imitator* hindlimbs sorted according to a* color value. Digital images of *R. imitator* hindlimbs corresponding to the red-highlighted region of interest in the inset photo (bottom right). The images have been sorted according to their CIELAB a* (green-red axis) value, with a* increasing from the top to bottom row as well as left to right; the top-left frog’s hindlimbs have the lowest (most green) value while the bottom-right frog’s hindlimbs have the highest (most red) value. Images labeled with a “B” and “S” indicate frogs sampled from the banded and striped morph populations respectively, and all other frogs were sampled from the admixture zone.

**Fig. S4.**
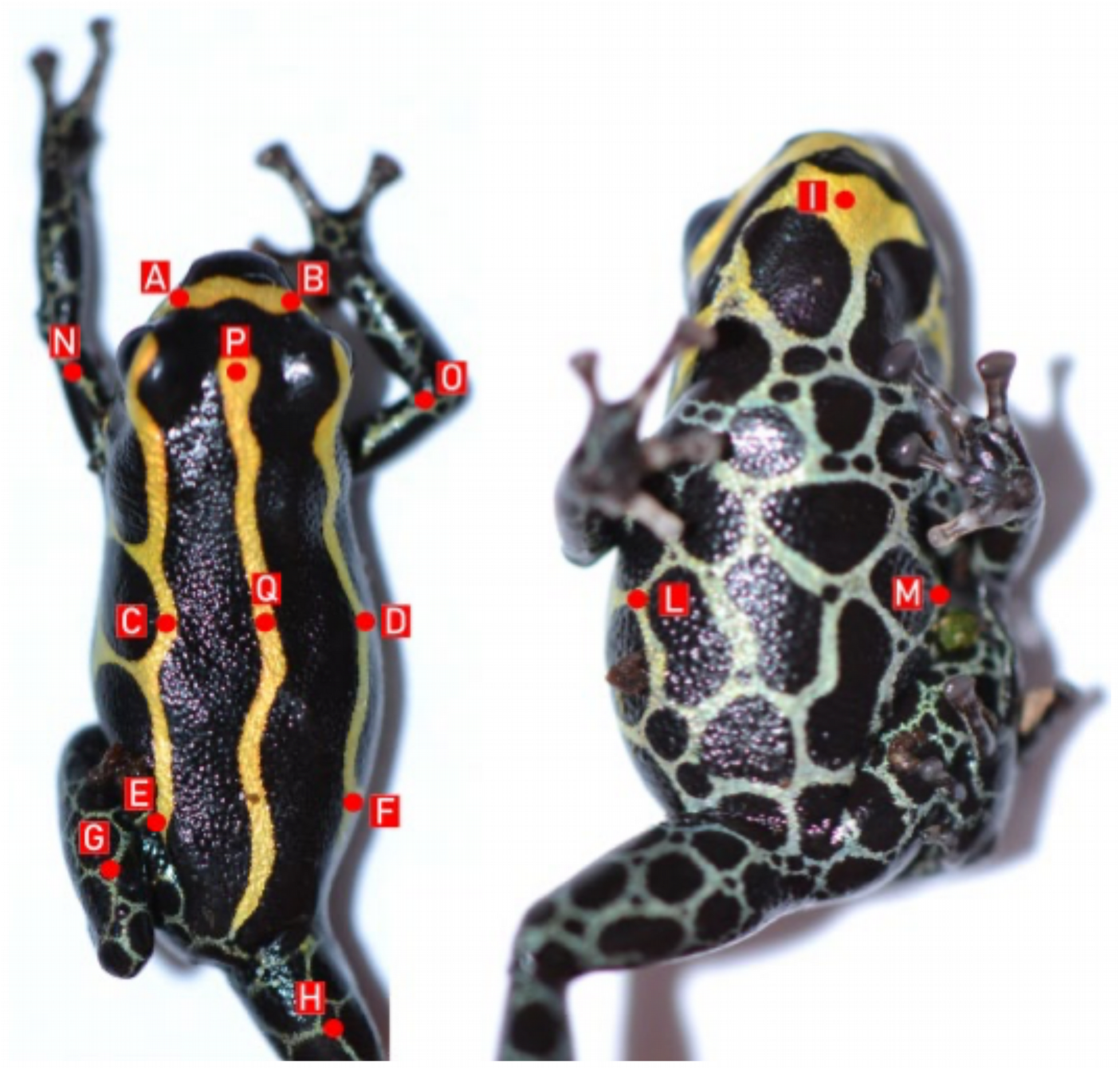
Positions on the body used for spectral reflectance measurements. Red, alphabetically-labeled points mark the locations for which spectral reflectance was measured on captive-reared frogs used in the genetic linkage mapping. Reflectance was always measured from a non-black point on each frog as near to the displayed points as possible.

**Fig. S5.**
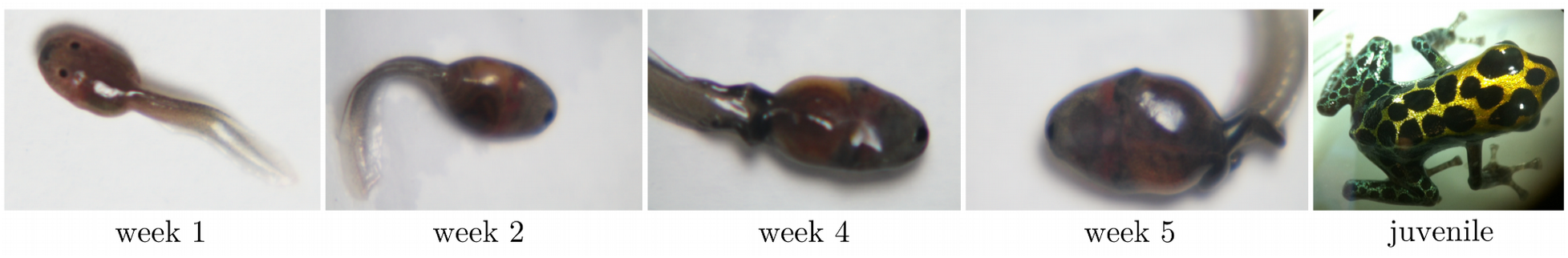
Stages of development used for *Ranitomeya imitator* transcriptome sequencing. Photos of five *R. imitator* individuals taken at the different time points throughout development for which we sequenced transcriptomes for exon capture probe design.

**Fig. S6.**
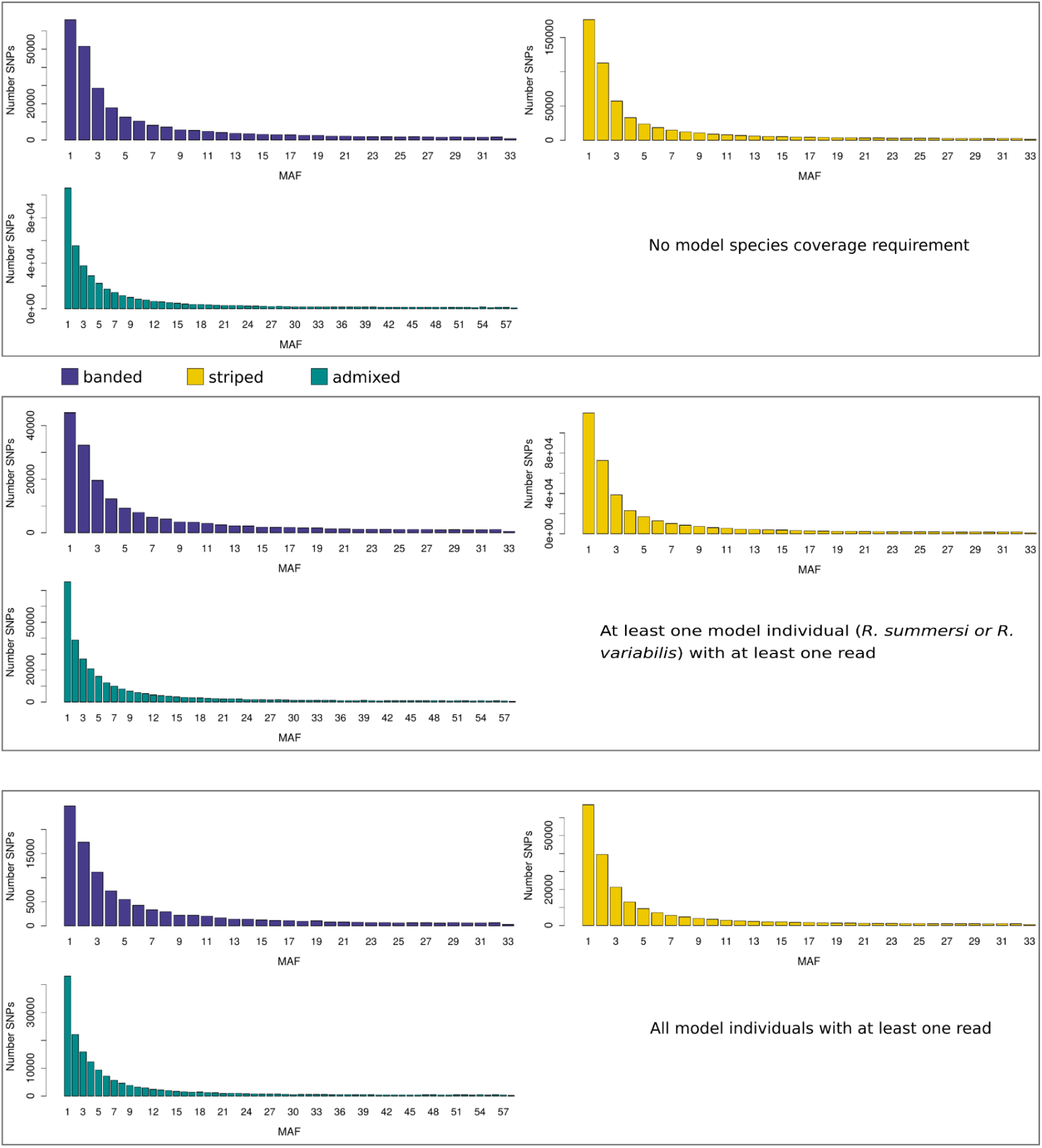
Site frequency spectrum for the banded, striped, and admixed *R. imitator* samples under different model species sequencing coverage requirements. Different depth of coverage criteria were used to obtain three subsets of sites, for which the respective SFS are shown in the top, middle, and bottom panel for each *R. imitator* population. The degree of representation among model species (*R. summersi* and *R. variabilis*) required for retained sites increases from the top to bottom panels at the expense of fewer SNPs among *R. imitator*. Higher representation among model species is important for interspecific comparisons while higher SNP density is conducive for intraspecific *R. imitator* analysis such as GWAS.

**Fig. S7.**
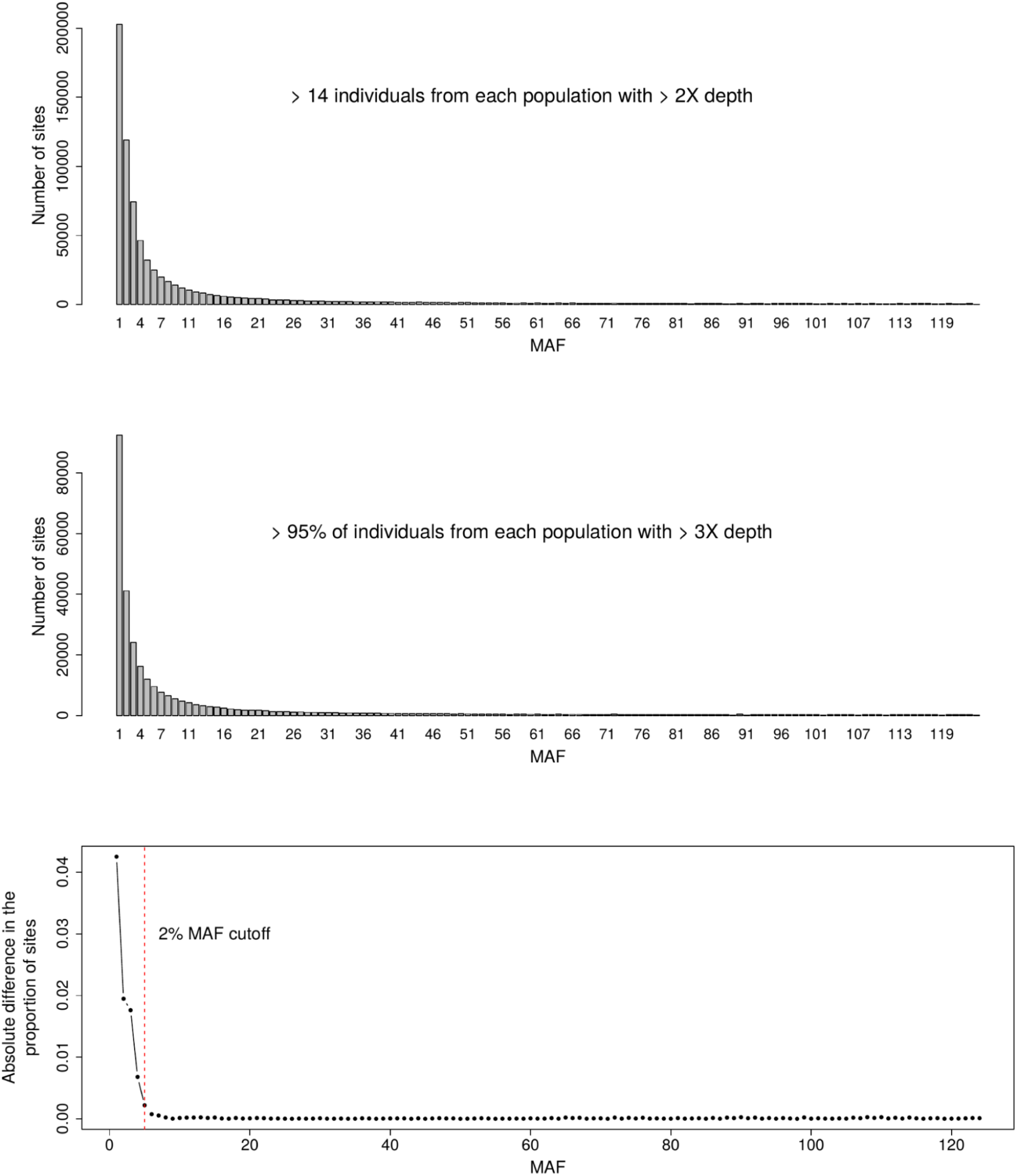
Effect of depth filters on the *R. imitator* site frequency spectrum. The proportion of sites belonging to different minor allele frequency classes below 2% was sensitive to different depth of coverage criteria for the *R. imitator* sample (bottom panel). The middle panel shows the *R. imitator* SFS estimated using stricter sequencing depth criteria compared to the top panel, leading to a higher proportion of singletons, but lower doubletons and tripletons for example. In order to ensure that inferences were robust to differences in how data were filtered we removed sites with MAF below 2% in *R. imitator* from all analyses.

**Fig. S8.**
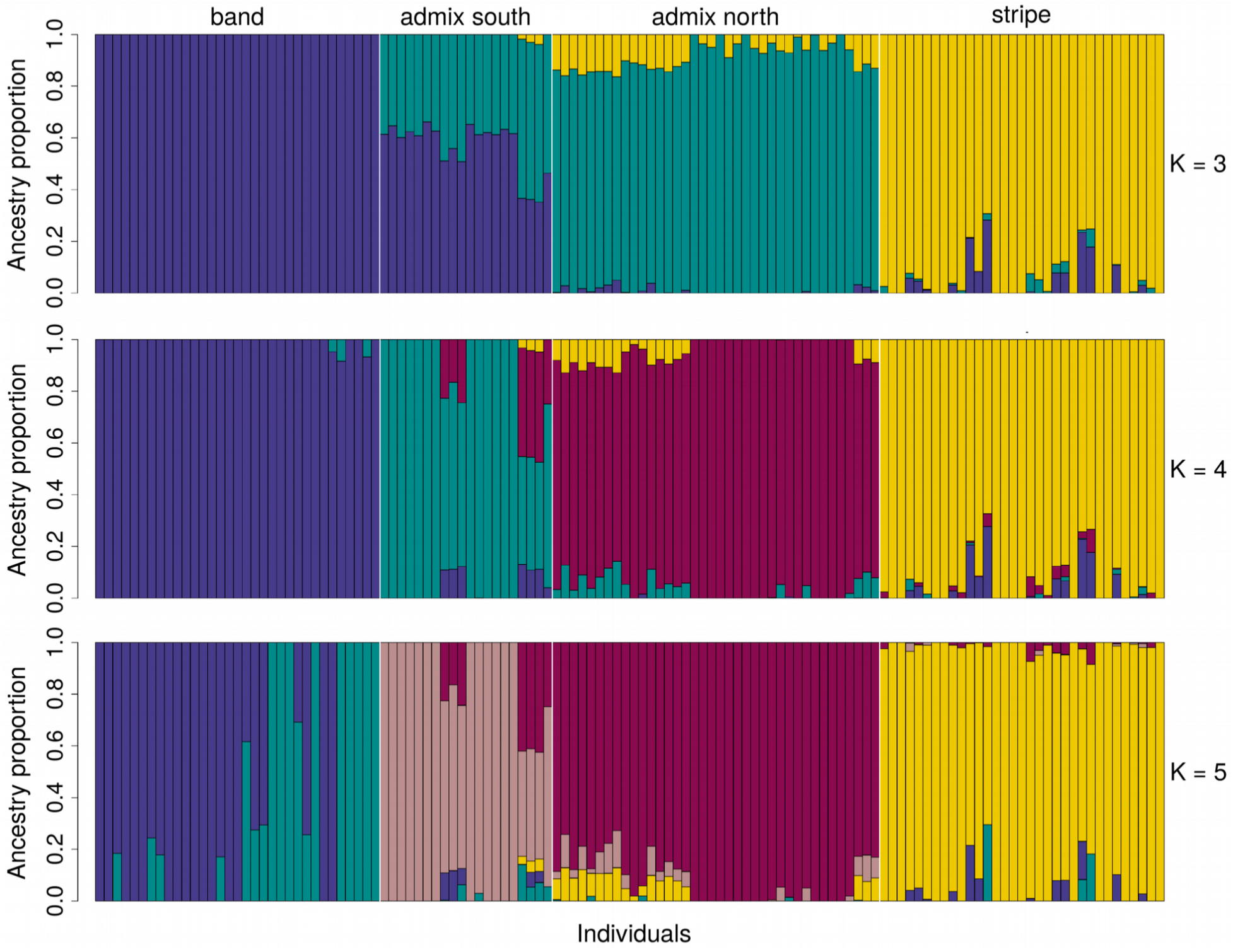
Admixture analysis based on 3-5 ancestral lineages. Proportions of distinct ancestries (different colors) for the 124 *R. imitator* individuals in our sample (bars) inferred using 3-5 ancestral lineages (K) with NGSadmix. Individuals have been grouped by population, which are delineated with vertical white bars. Individuals from the “admix south” and “admix north” populations were sampled south and north of the Huallaga River, respectively (Fig. 1).

**Fig. S9.**
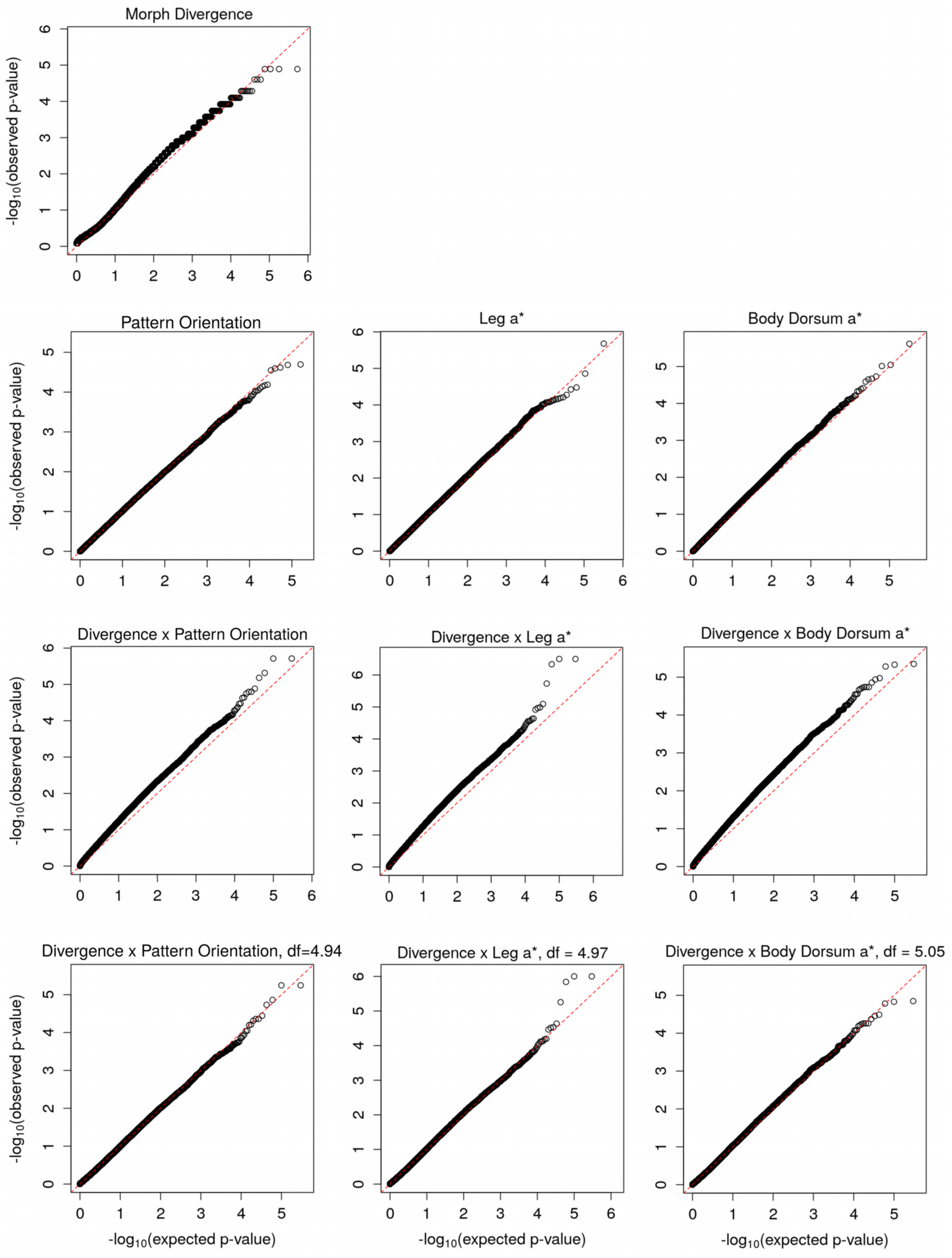
Comparisons of the observed versus expected distribution of p-values for genome-wide association tests. Quantile-quantile (QQ) plots of the observed versus expected distribution of p-values for the divergence test between banded and striped morphs (top row) and the association test using general linear models to relate genotypes to pattern orientation, hindlimb a*, and body dorsum a* (second-from-top row). The third-from-top row shows p-value QQ-plots for the X^2^ statistic from combining p-values from the two aforementioned tests using Fisher’s method and using the theoretical *X*^2^_4_ null distribution. The bottom row shows p-value QQ-plots for the combined p-value X^2^ statistic using a *X*^2^*_df_* null distribution where we fit the degrees of freedom (*df*) to the empirical X^2^ distribution.

**Fig. S10.**
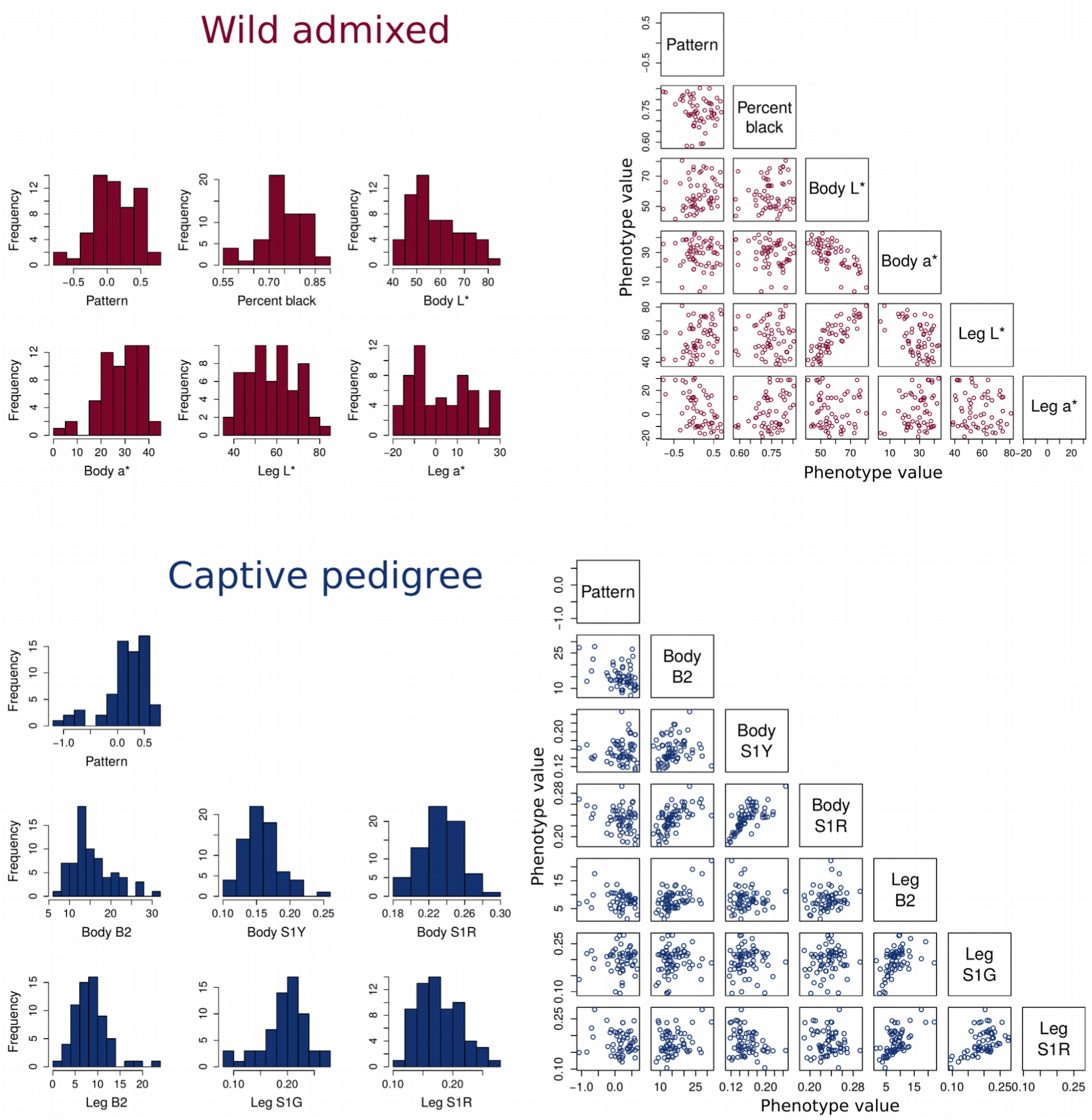
Distribution of morph-related traits in captive and wild *R. imitator*. Histograms show the distribution of phenotypic values for color morph traits while scatterplots show relationships between them among wild, admixed individuals used for genetic association mapping (top, maroon) and captive pedigree individuals used for linkage mapping (bottom, blue). For the wild population individuals, we measured pattern orientation (orientation of pattern along the body dorsum), percentage of the body dorsum that was black, and the value of the non-black portions of the body and hindlimb (“leg”) dorsal surfaces along the lightness (L*) and red-green (a*) color axes of the CIELAB color space. For the captive pedigree individuals, we measured pattern orientation as well as the brightness and yellow (S1Y), red (S1R), and green (S1G) wavelength reflectance for the non-black portions of the body and/or hindlimb dorsal surfaces.

**Table S1.**
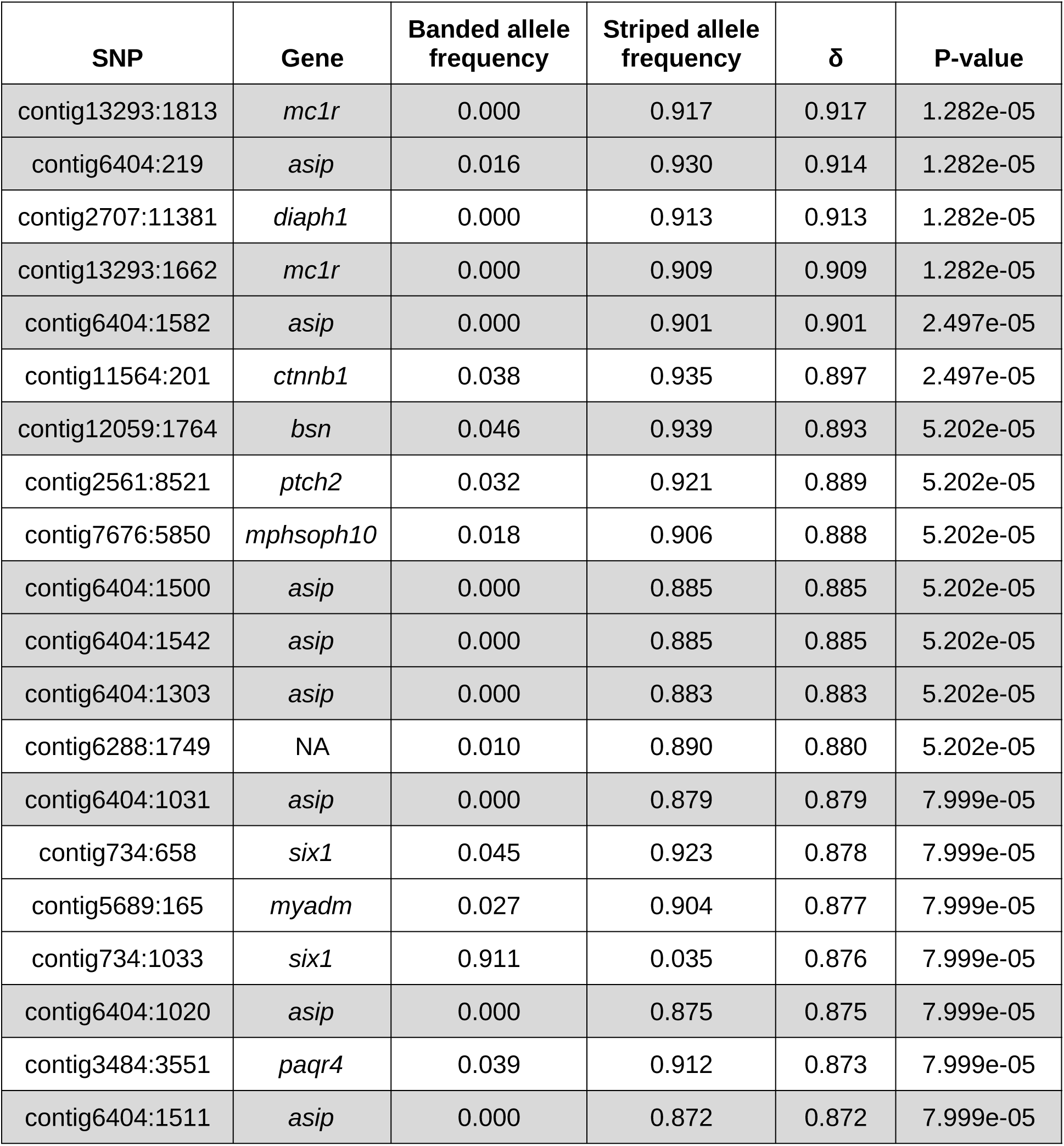
The top 20 most divergent SNPs between the banded and striped R. imitator morphs in descending order of absolute allele frequency difference, δ. P-values correspond to the probability, given our sample sizes, of observing allele frequency differences larger than δ by chance under a Balding-Nichols model of pure drift. SNPs in genes deemed as candidates for influencing coloration are highlighted in gray.

**Table S2.**
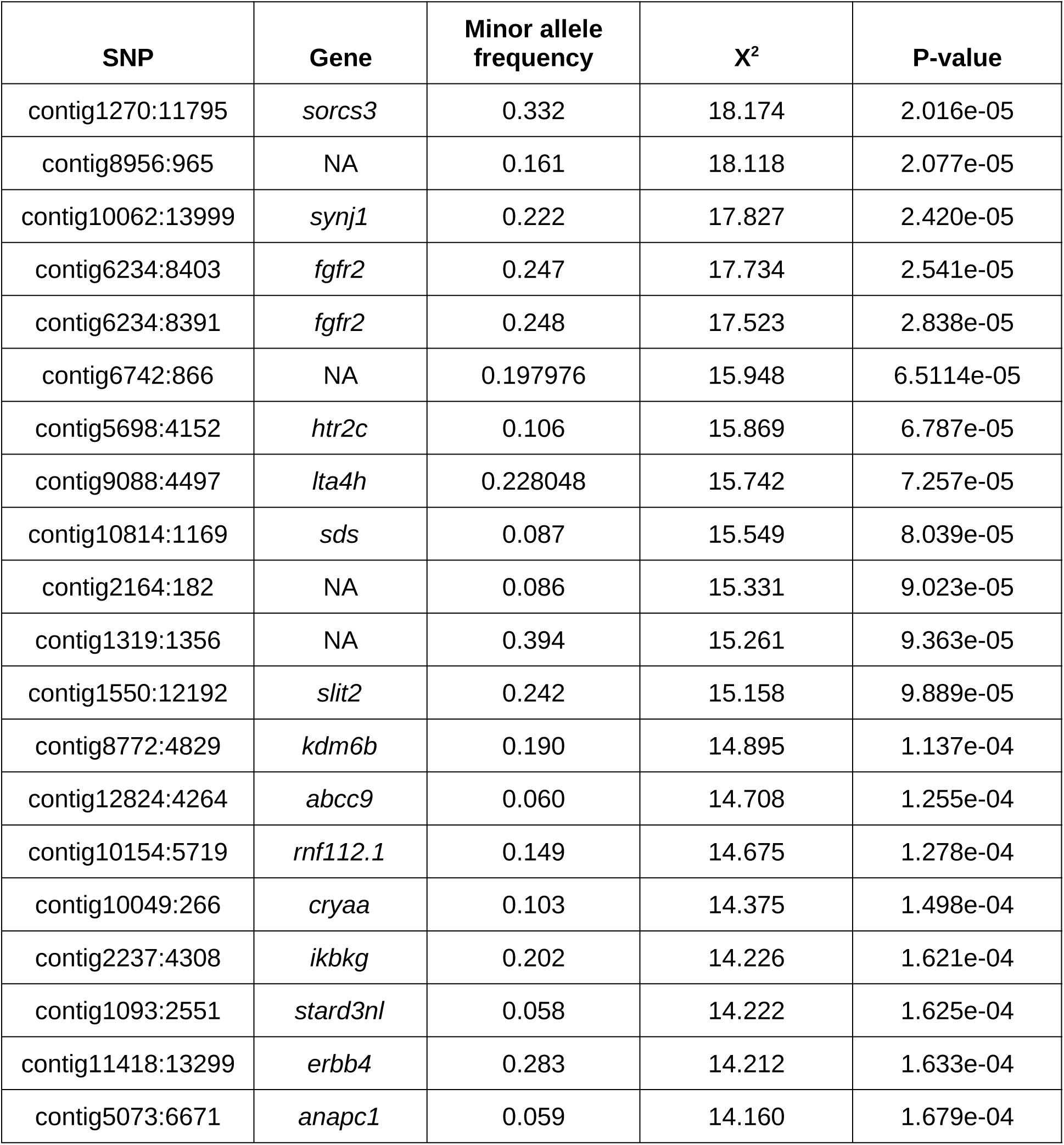
The top 20 SNPs with the highest genotypic association with pattern orientation in the wild, admixed population of *R. imitator*. SNPs are listed in descending order of the association likelihood ratio test statistic, X^2^, along with corresponding p-values.

| SNP | Gene | Minor allele frequency | $X^2$ | P-value |
| --- | --- | --- | --- | --- |
| contig1270:11795 | *sorcs3* | 0.332 | 18.174 | 2.016e-05 |
| contig8956:965 | NA | 0.161 | 18.118 | 2.077e-05 |
| contig10062:13999 | *synj1* | 0.222 | 17.827 | 2.420e-05 |
| contig6234:8403 | *fgfr2* | 0.247 | 17.734 | 2.541e-05 |
| contig6234:8391 | *fgfr2* | 0.248 | 17.523 | 2.838e-05 |
| contig6742:866 | NA | 0.197976 | 15.948 | 6.5114e-05 |
| contig5698:4152 | *htr2c* | 0.106 | 15.869 | 6.787e-05 |
| contig9088:4497 | *lta4h* | 0.228048 | 15.742 | 7.257e-05 |
| contig10814:1169 | *sds* | 0.087 | 15.549 | 8.039e-05 |
| contig2164:182 | NA | 0.086 | 15.331 | 9.023e-05 |
| contig1319:1356 | NA | 0.394 | 15.261 | 9.363e-05 |
| contig1550:12192 | *slit2* | 0.242 | 15.158 | 9.889e-05 |
| contig8772:4829 | *kdm6b* | 0.190 | 14.895 | 1.137e-04 |
| contig12824:4264 | *abcc9* | 0.060 | 14.708 | 1.255e-04 |
| contig10154:5719 | *rnf112.1* | 0.149 | 14.675 | 1.278e-04 |
| contig10049:266 | *cryaa* | 0.103 | 14.375 | 1.498e-04 |
| contig2237:4308 | *ikbkg* | 0.202 | 14.226 | 1.621e-04 |
| contig1093:2551 | *stard3nl* | 0.058 | 14.222 | 1.625e-04 |
| contig11418:13299 | *erbb4* | 0.283 | 14.212 | 1.633e-04 |
| contig5073:6671 | *anapc1* | 0.059 | 14.160 | 1.679e-04 |

**Table S3.**
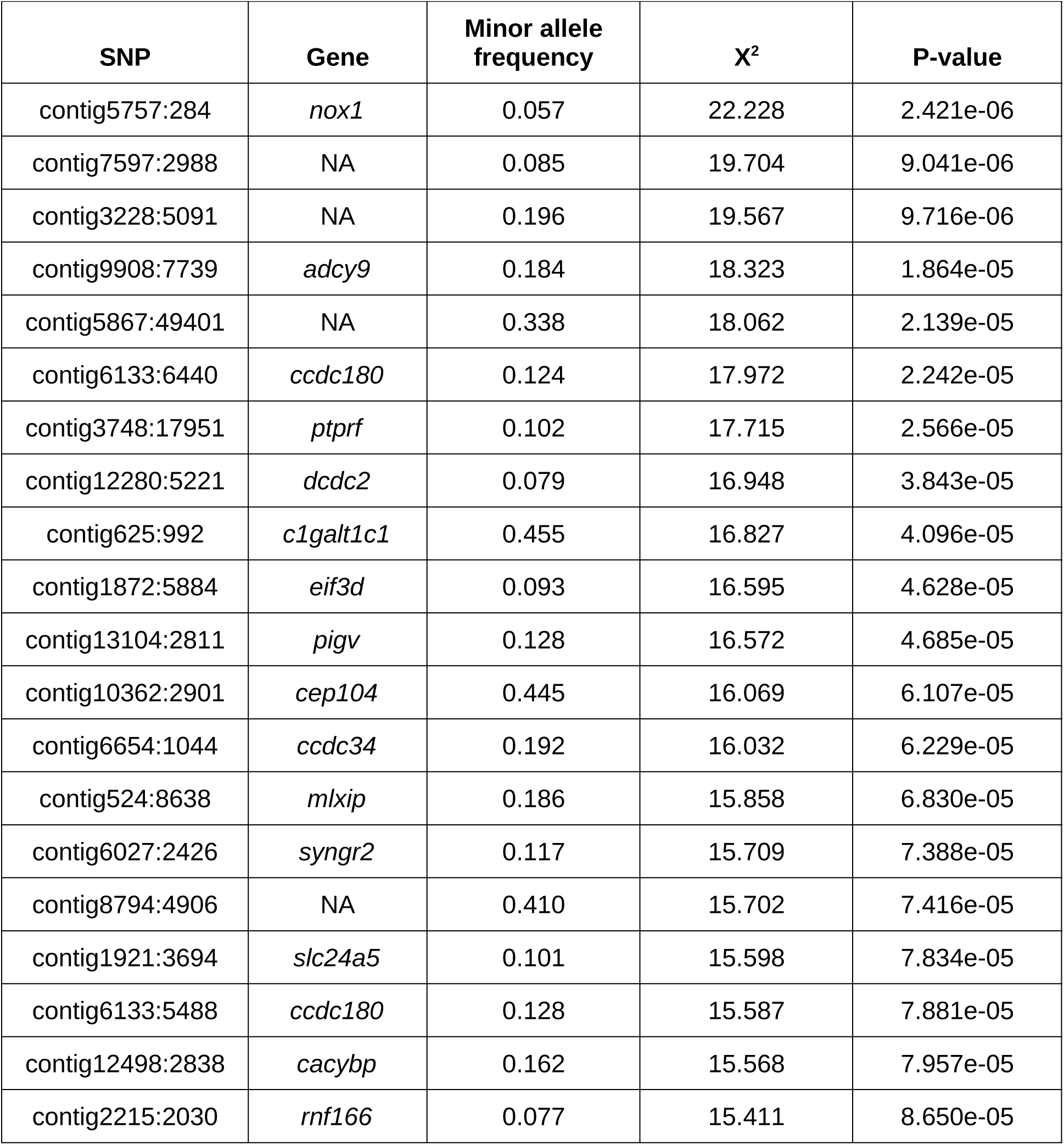
The top 20 SNPs with the highest genotypic association with dorsal body color in the wild, admixed population of R. imitator. SNPs are listed in descending order of the association likelihood ratio test statistic, X^2^, along with corresponding p-values.

**Table S4.**
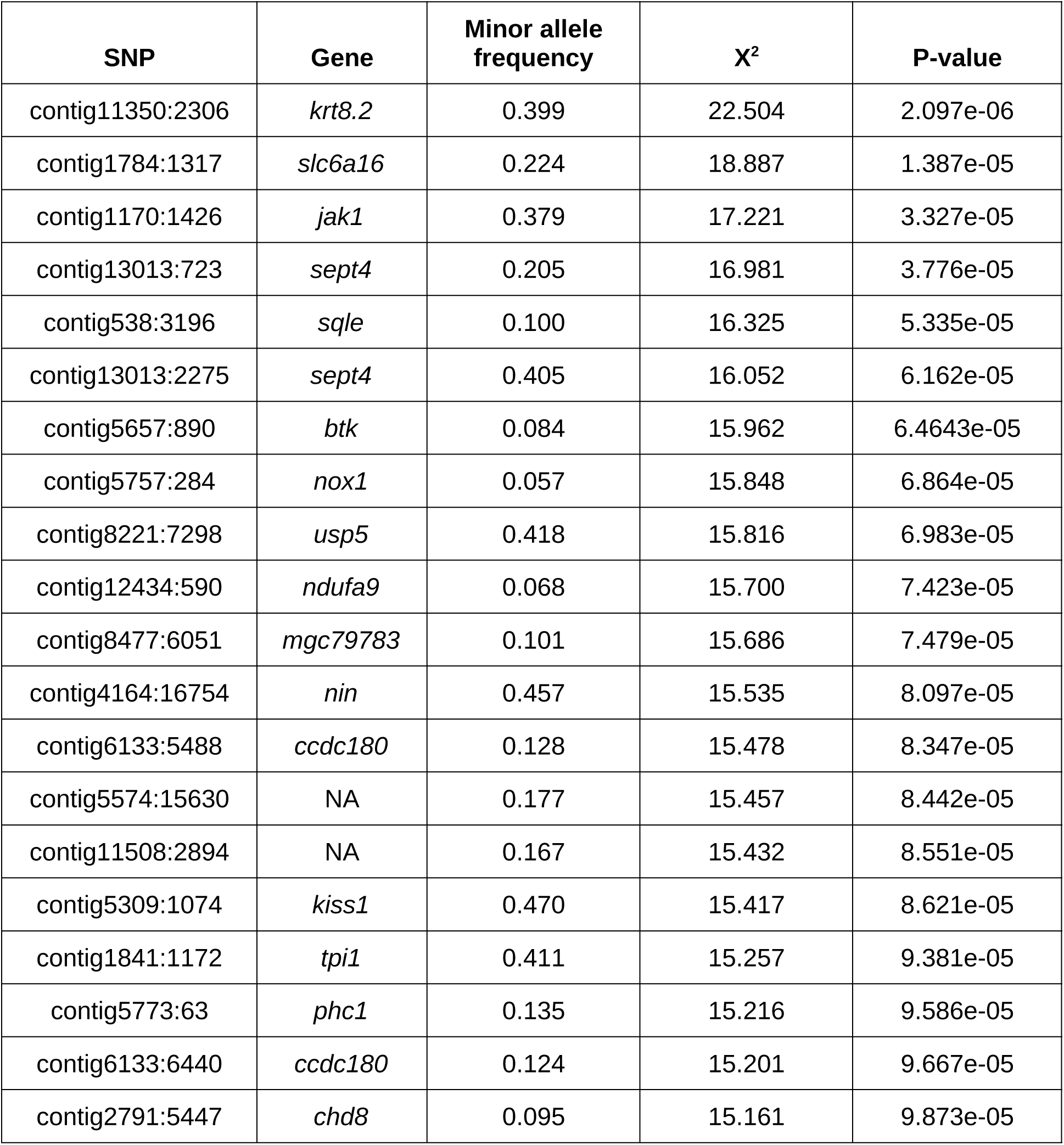
The top 20 SNPs with the highest genotypic association with hindlimb color in the wild, admixed population of *R. imitator*. SNPs are listed in descending order of the association likelihood ratio test statistic, X^2^, along with corresponding p-values.

**Table S5.** The top 20 SNPs with the most evidence for influencing pattern orientation based on combined p-values from the test of excessive allele frequency divergence between the striped and banded morphs of R. imitator and the association test for pattern orientation in the admixed population. SNPs are sorted in descending order of Fisher’s combined p-value statistic, *X*^2^*_combined_*, with corresponding p-values after genomic control. The genome-wide quantiles for the absolute allele frequency difference, δ, between the striped and banded morphs and the likelihood ratio statistic, X^2^*_admix_*, for the test of an association with pattern orientation in the admixed population are also shown for each SNP. SNPs in candidate pattern genes are highlighted in gray.

| SNP | Gene | $X^2_{combined}$ | P-value | $\delta$ quantile | $X^2_{admix}$ quantile |
| --- | --- | --- | --- | --- | --- |
| contig13293:1813 | *mc1r* | 31.97478 | 5.655e-06 | 1.0000000 | 0.9913467 |
| contig13293:1662 | *mc1r* | 31.97478 | 5.655e-06 | 0.9999888 | 0.9913467 |
| contig12059:1764 | *bsn* | 29.36860 | 1.851e-05 | 0.9999775 | 0.9922763 |
| contig11798:9830 | *clip1* | 27.89937 | 3.597e-05 | 0.9990148 | 0.9990704 |
| contig1270:11795 | *sorcs3* | 27.48134 | 4.342e-05 | 0.9352374 | 1.0000000 |
| contig734:1033 | *six1* | 27.46881 | 4.367e-05 | 0.9999401 | 0.9865427 |
| contig4416:930 | *decr2* | 26.70407 | 6.158e-05 | 0.9995767 | 0.9943476 |
| contig10062:13999 | *synj1* | 26.62754 | 6.373e-05 | 0.926715 | 0.9999875 |
| contig6742:866 | NA | 25.87821 | 8.916e-05 | 0.9515741 | 0.9999688 |
| contig8326:5212 | NA | 25.82889 | 9.115e-05 | 0.9983217 | 0.9977602 |
| contig6404:1582 | *asip* | 25.36693 | 1.120e-04 | 0.9999850 | 0.8723087 |
| contig7105:677 | *fam178a* | 25.15061 | 1.234e-04 | 0.9975163 | 0.9975419 |
| contig6769:405 | NA | 24.92548 | 1.364e-04 | 0.9999063 | 0.9667218 |
| contig9196:5283 | *rhobtb1* | 24.88606 | 1.388e-04 | 0.9998614 | 0.9661041 |
| contig3681:405 | *aifm1* | 24.42162 | 1.707e-04 | 0.9982431 | 0.9953208 |
| contig6404:219 | *asip* | 24.27235 | 1.824e-04 | 0.9999963 | 0.5774028 |
| contig6931:14551 | *fancd2* | 24.22644 | 1.867e-04 | 0.9997528 | 0.9692548 |
| contig12985:4310 | NA | 24.09677 | 1.972e-04 | 0.9994381 | 0.9846711 |
| contig6989:5080 | *smarca2* | 24.07623 | 1.990e-04 | 0.9996741 | 0.9776086 |
| contig2491:447 | *barx1* | 24.05234 | 2.011e-04 | 0.9658205 | 0.9998004 |

**Table S6.**
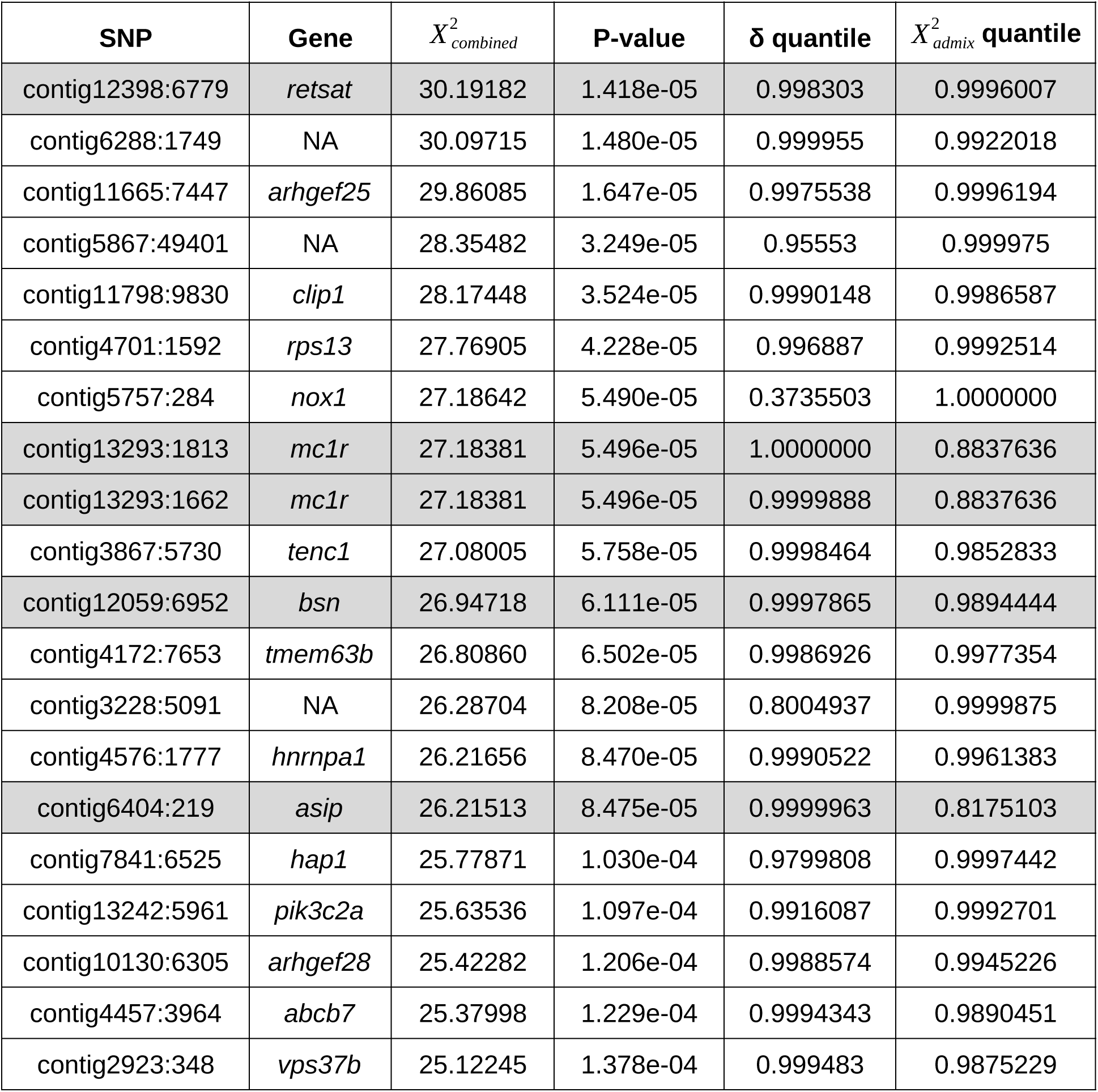
The top 20 SNPs with the most evidence for influencing dorsal body color based on combined p-values from the test of excessive allele frequency divergence between the striped and banded morphs of *R. imitator* and the association test for dorsal body color in the admixed population. SNPs are sorted in descending order of Fisher’s combined p-value statistic, *X*^2^*_combined_*, with corresponding p-values after genomic control. The genome-wide quantiles for the absolute allele frequency difference, δ, between the striped and banded morphs and the likelihood ratio statistic, *X*^2^*_admix_*, for the test of an association with dorsal body color in the admixed population are also shown for each SNP. SNPs in candidate color genes are highlighted in gray.

**Table S7.**
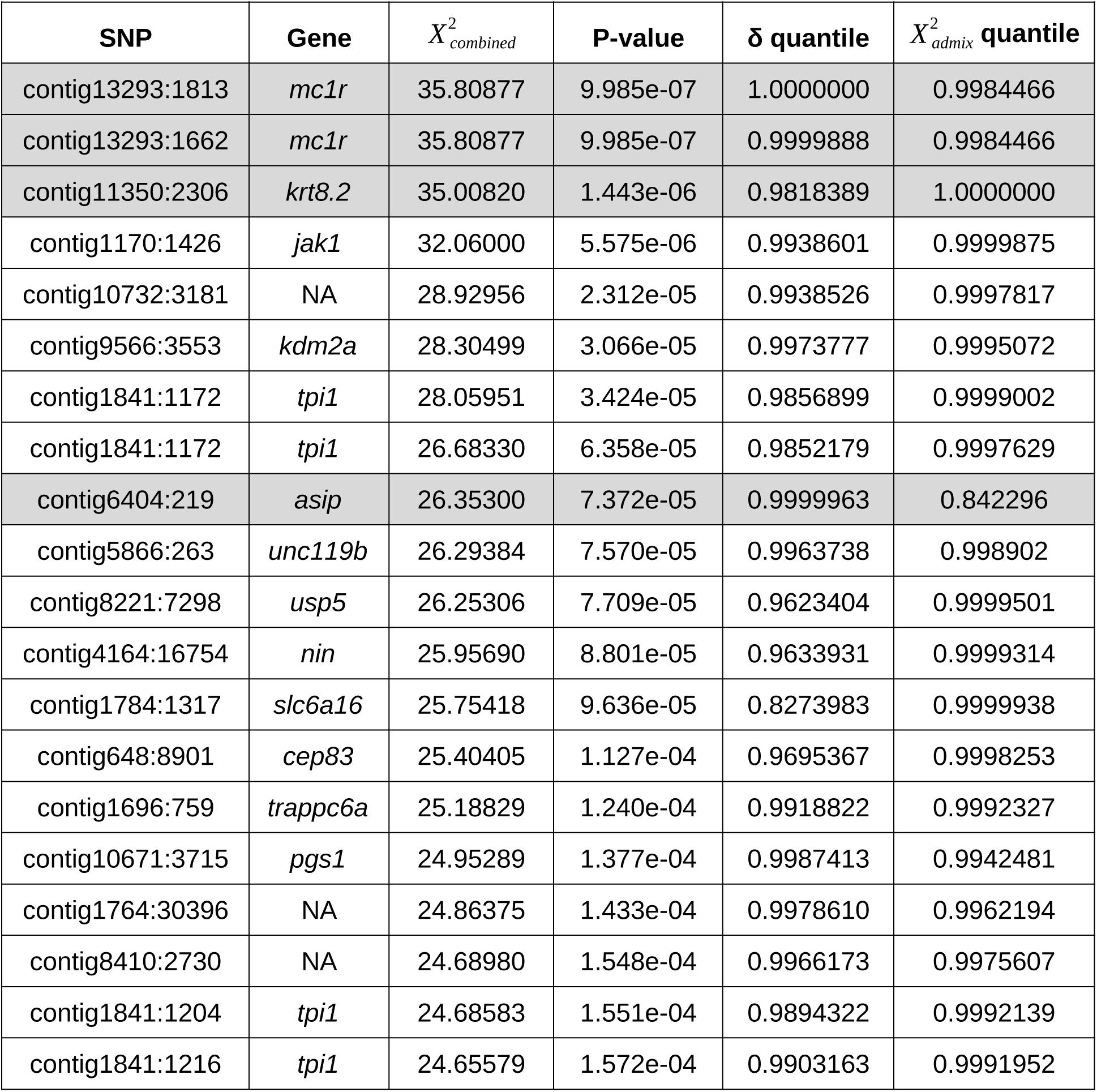
The top 20 SNPs with the most evidence for influencing hindlimb color based on combined p-values from the test of excessive allele frequency divergence between the striped and banded morphs of R. imitator and the association test for hindlimb color in the admixed population. SNPs are sorted in descending order of the combined p-value statistic, X^2^*_combined_*, with corresponding p-values after genomic control. The genome-wide quantiles for the absolute allele frequency difference, δ, between the striped and banded morphs and the likelihood ratio statistic, X^2^*_admix_*, for the test of an association with hindlimb color in the admixed population are also shown for each SNP. SNPs in candidate color genes are highlighted in gray.

| SNP | Gene | $X^2_{combined}$ | P-value | $\delta$ quantile | $X^2_{admix}$ quantile |
| --- | --- | --- | --- | --- | --- |
| contig13293:1813 | *mc1r* | 35.80877 | 9.985e-07 | 1.0000000 | 0.9984466 |
| contig13293:1662 | *mc1r* | 35.80877 | 9.985e-07 | 0.9999888 | 0.9984466 |
| contig11350:2306 | *krt8.2* | 35.00820 | 1.443e-06 | 0.9818389 | 1.0000000 |
| contig1170:1426 | *jak1* | 32.06000 | 5.575e-06 | 0.9938601 | 0.9999875 |
| contig10732:3181 | NA | 28.92956 | 2.312e-05 | 0.9938526 | 0.9997817 |
| contig9566:3553 | *kdm2a* | 28.30499 | 3.066e-05 | 0.9973777 | 0.9995072 |
| contig1841:1172 | *tpi1* | 28.05951 | 3.424e-05 | 0.9856899 | 0.9999002 |
| contig1841:1172 | *tpi1* | 26.68330 | 6.358e-05 | 0.9852179 | 0.9997629 |
| contig6404:219 | *asip* | 26.35300 | 7.372e-05 | 0.9999963 | 0.842296 |
| contig5866:263 | *unc119b* | 26.29384 | 7.570e-05 | 0.9963738 | 0.998902 |
| contig8221:7298 | *usp5* | 26.25306 | 7.709e-05 | 0.9623404 | 0.9999501 |
| contig4164:16754 | *nin* | 25.95690 | 8.801e-05 | 0.9633931 | 0.9999314 |
| contig1784:1317 | *slc6a16* | 25.75418 | 9.636e-05 | 0.8273983 | 0.9999938 |
| contig648:8901 | *cep83* | 25.40405 | 1.127e-04 | 0.9695367 | 0.9998253 |
| contig1696:759 | *trappc6a* | 25.18829 | 1.240e-04 | 0.9918822 | 0.9992327 |
| contig10671:3715 | *pgs1* | 24.95289 | 1.377e-04 | 0.9987413 | 0.9942481 |
| contig1764:30396 | NA | 24.86375 | 1.433e-04 | 0.9978610 | 0.9962194 |
| contig8410:2730 | NA | 24.68980 | 1.548e-04 | 0.9966173 | 0.9975607 |
| contig1841:1204 | *tpi1* | 24.68583 | 1.551e-04 | 0.9894322 | 0.9992139 |
| contig1841:1216 | *tpi1* | 24.65579 | 1.572e-04 | 0.9903163 | 0.9991952 |

**Table S8.** Phenotypic variance explained by SNPs in candidate genes after accounting for genome-wide ancestry and technical covariates.

| **Phenotype** | **SNP** | **Variance explained (%)** |
| --- | --- | --- |
| pattern orientation | *mc1r*:1813 | 9.0 |
| pattern orientation | *asip*:1582 | 3.8 |
| pattern orientation | *bsn*:1764 | 9.0 |
| body color | *mc1r*:1813 | 2.5 |
| body color | *asip*:219 | 1.8 |
| body color | *bsn*:6952 | 6.5 |
| body color | *retsat*:6779 | 12.0 |
| hindlimb color | *mc1r*:1813 | 8.3 |
| hindlimb color | *asip*:219 | 1.2 |
| hindlimb color | *krt8.2*:2306 | 17.6 |

**Table S9.** Logarithm of the odds (LOD) for an association between candidate gene SNPs and color phenotypes in the R. imtitator pedigree. Significant associations with FDR below 10% are shaded gray.

| **Gene** | **Phenotype** | **LOD score** | **q-value** |
| --- | --- | --- | --- |
| *mc1r* | pattern orientation | 2.52 | 0.003 |
| *mc1r* | body brightness (B2) | 0.28 | 0.350 |
| *mc1r* | body chroma (S1Y) | 0.21 | 0.420 |
| *mc1r* | hindlimb brightness (B2) | 1.13 | 0.064 |
| *asip* | pattern orientation | 2.54 | 0.003 |
| *asip* | body brightness (B2) | 2.47 | 0.003 |
| *asip* | body chroma (S1Y) | 0.17 | 0.432 |
| *asip* | hindlimb brightness (B2) | 0.62 | 0.182 |
| *asip* | hindlimb chroma (S1G) | 0.00 | 0.950 |
| *retsat* | body brightness (B2) | 0.33 | 0.342 |
| *retsat* | body chroma (S1Y) | 3.49 | 0.001 |
| *bsn* | pattern orientation | 0.93 | 0.091 |
| *bsn* | body brightness (B2) | 0.09 | 0.571 |
| *bsn* | body chroma (S1Y) | 0.48 | 0.238 |

**Table S10.** D-statistic test for genome-wide introgression between the model species and *R. imitator*. No Significant differences in the number of ABBA and BABA allele configurations among banded *R. imitator* (H1), striped *R. imitator* (H2), *R. variabilis* (H3), and *R. summersi* (outgroup) were identified using Z-tests on the genome-wide, mean D-statistic and D-statistic corrected for bias (JK-D) from jackknife resampling of “nBlocks” genomic blocks. V(JK-D) is the estimate of the variance for D across blocks used to calculate the Z-statistic.

| D | JK-D | V(JK-D) | Z | P-value | ABBA | BABA | nBlocks |
| --- | --- | --- | --- | --- | --- | --- | --- |
| 0.00246 | 0.00246 | 0.00002 | 0.58527 | 0.55837 | 637.16901 | 634.0413 | 11247 |

**Table S11.** Observed and expected counts of segregating sites within R. imitator and fixed differences from the model species, *R. summersi* and *R. variabilis*, at candidate gene exons containing color-associated SNPs. The ratio of the observed number of *R. imitator* segregating sites (S_O_) to the observed number of fixed differences from the model species (D_o_) was compared to the ratio of the expected number of segregating sites (S_E_) to the expected number of fixed differences (D_E_) under neutrality at the candidate exons using Hudson–Kreitman–Aguadé (HKA) tests. Exons for which the HKA test significantly rejects a model of neutral evolution at a 5% false discovery rate are shaded gray. Notice that a correction for genome-wide deviation from the theoretical X^2^_1_ distribution has been applied to the X^2^ values.

| Morph | Locus | $S_o$ | $S_E$ | $D_o$ | $D_E$ | $X^2$ | q-value |
| --- | --- | --- | --- | --- | --- | --- | --- |
| *R. imitator*<br>(banded and striped combined) | *mc1r* exon 2 | 7 | 13.19 | 37 | 30.81 | 6.25 | 0.032 |
|  | *asip* exon 1 | 2 | 1.50 | 3 | 3.50 | 4.03e-6 | 0.998 |
|  | *asip* exon 2 | 7 | 5.09 | 10 | 11.91 | 0.99 | 0.481 |
|  | *bsn* exon 2 | 15 | 30.27 | 86 | 70.73 | 18.35 | 1.101e-4 |
|  | *retsat* exon 11 | 2 | 6.29 | 19 | 14.71 | 5.82 | 0.032 |
|  | *krt8.2* exon 3 | 3 | 2.10 | 4 | 4.90 | 0.20 | 0.789 |
| Banded *R. imitator* | *mc1r* exon 2 | 0 | 9.02 | 42 | 32.98 | 22.44 | 1.304e-5 |
|  | *asip* exon 1 | 2 | 1.07 | 3 | 3.93 | 0.47 | 0.591 |
|  | *asip* exon 2 | 0 | 2.58 | 12 | 9.42 | 4.67 | 0.046 |
|  | *bsn* exon 2 | 10 | 20.83 | 87 | 76.17 | 14.28 | 4.717e-4 |
|  | *retsat* exon 11 | 0 | 4.51 | 21 | 16.49 | 9.94 | 0.003 |
|  | *krt8.2* exon 3 | 2 | 1.29 | 4 | 4.71 | 0.10 | 0.756 |
| Striped *R.* | *mc1r* | 7 | 12.37 | 37 | 31.63 | 4.46 | 0.069 |
| *imitator* | exon 2 |  |  |  |  |  |  |
|  | *asip*<br>exon 1 | 2 | 1.41 | 3 | 3.59 | 0.01 | 0.903 |
|  | *asip*<br>exon 2 | 7 | 4.78 | 10 | 12.22 | 1.44 | 0.345 |
|  | *bsn*<br>exon 2 | 14 | 28.11 | 86 | 71.87 | 15.34 | 5.394e-4 |
|  | *retsat*<br>exon 11 | 2 | 5.90 | 19 | 15.10 | 4.57 | 0.069 |
|  | *krt8.2*<br>exon 3 | 2 | 1.69 | 4 | 4.31 | 0.05 | 0.903 |

**Table S12.**
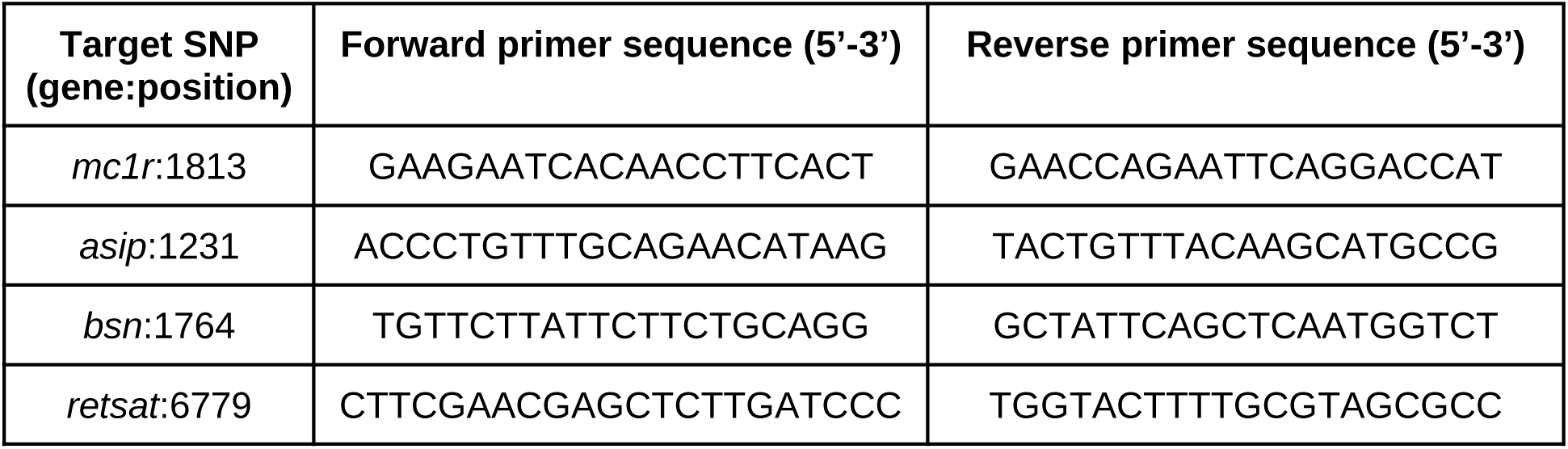
Sequencing primers for genotyping candidate gene SNPs in the *R. imitator* captive pedigree.

## Notes

### Competing Interest Statement

The authors have declared no competing interest.

